# Dorsal and ventral hippocampus engage opposing networks in the nucleus accumbens

**DOI:** 10.1101/604116

**Authors:** Marielena Sosa, Hannah R. Joo, Loren M. Frank

**Affiliations:** Neuroscience Graduate Program, Kavli Institute for Fundamental Neuroscience and Department of Physiology, University of California San Francisco, CA 94158, USA.; Howard Hughes Medical Institute, San Francisco, CA 94158, USA.

## Abstract

Memories of positive experiences require the brain to link places, events, and reward outcomes. Neural processing underlying the association of spatial experiences with reward is thought to depend on interactions between the hippocampus and the nucleus accumbens (NAc)^1–9^. Hippocampal projections to the NAc arise from both the ventral hippocampus (vH) and the dorsal hippocampus (dH)^6–12^, and studies using optogenetic interventions have demonstrated that either vH^5, 6^ or dH^7^ input to the NAc can support behaviors dependent on spatial-reward associations. It remains unclear, however, whether dH, vH, or both coordinate memory processing of spatial-reward information in the hippocampal-NAc circuit under normal conditions. Times of memory reactivation within and outside the hippocampus are marked by hippocampal sharp-wave ripples (SWRs)^13–19^, discrete events which facilitate investigation of inter-regional information processing. It is unknown whether dH and vH SWRs act in concert or separately to engage NAc neuronal networks, and whether either dH or vH SWRs are preferentially linked to spatial-reward representations. Here we show that dH and vH SWRs occur asynchronously in the awake state and that NAc spatial-reward representations are selectively activated during dH SWRs. We performed simultaneous extracellular recordings in the dH, vH, and NAc of rats learning and performing an appetitive spatial task and during sleep. We found that individual NAc neurons activated during SWRs from one subdivision of the hippocampus were typically suppressed or unmodulated during SWRs from the other. NAc neurons activated during dH versus vH SWRs showed markedly different task-related firing patterns. Only dH SWR-activated neurons were tuned to similarities across spatial paths and past reward, indicating a specialization for the dH-NAc, but not vH-NAc, network in linking reward to discrete spatial paths. These temporally and anatomically separable hippocampal-NAc interactions suggest that dH and vH coordinate opposing channels of mnemonic processing in the NAc.

## Text

Identifying the nature of coordination between the dH-NAc and vH-NAc pathways requires a simultaneous survey of all three regions. We therefore recorded from dH, vH, and NAc using chronically implanted tetrode arrays in rats (Fig. 1a, Extended Data Fig. 1), in the context of both a dynamic spatial memory task and interleaved sleep periods. We utilized a “Multiple-W” task^19^ in which a rat must first learn which three of six maze arms are reward locations and alternate between them, then must transfer the alternation rule to a new set of reward locations (Fig. 1b,c, Extended Data Fig. 2). This task requires reward-based spatial learning and is known to engage dH SWRs^18, 19^, but when vH SWRs occur with respect to awake behavior is unknown.

**Figure 1.**
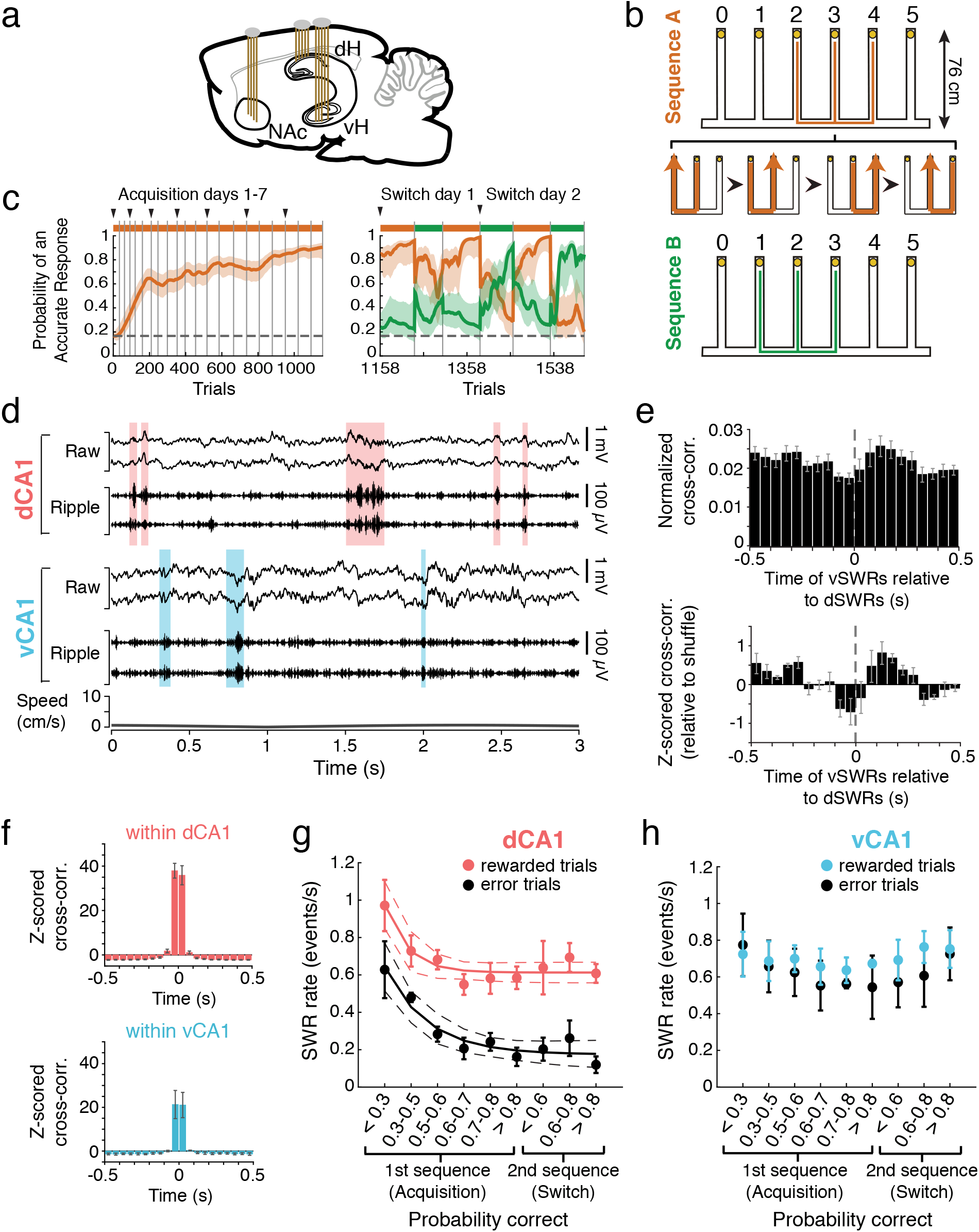
Awake dorsal and ventral hippocampal SWRs occur asynchronously. **a**, Sagittal schematic of tetrodes targeting NAc, dH, and vH in the rat brain. **b**, Schematic of the Multiple-W task. Yellow circles indicate visually identical reward wells. The animal was required to alternate visits from the center well of the “W” to the outer two wells to receive a liquid food reward on each correct well visit. Expanded section depicts four consecutive correct trials. Animals acquired one sequence (either A or B, counterbalanced across animals) to ∼80% correct (“Acquisition” days) before the second sequence was introduced on the first “Switch” day. Sequences were thereafter switched across maze epochs (behavior sessions separated by sleep epochs in a separate rest box) for the remainder of the experiment (Extended Data Fig. 2). **c**, Example behavior (Rat 5) expressed as the probability that the rat is making an accurate choice on each trial according to Sequence A (orange) or B (green) (see Online Methods). Solid line indicates the mode of the probability distribution, shaded region indicates the 90% confidence interval. Colored bars above the plot indicate the rewarded sequence, grey vertical lines mark epoch boundaries, black triangles mark the start of each day, and the horizontal dotted line indicates chance performance (0.167). For clarity, only 2 full switch days are shown on the right (see Extended Data Fig. 2 for the complete behavior). **d**, Example dSWRs and vSWRs during awake immobility at a reward well (Rat 4). The raw (1-400 Hz) and ripple-filtered (150-250 Hz) local field potential is shown for two tetrodes each in dorsal CA1 (dCA1) and ventral CA1 (vCA1). Shaded regions highlight detected dSWRs (pink) and vSWRs (blue). Bottom plot depicts the speed of the animal. **e**, Mean cross-correlation histogram (CCH) between onset times of dSWRs and vSWRs across animals (n=5 rats). Top: CCH normalized by the number of dSWRs. The y-axis signifies the fraction of dSWRs that have a vSWR occurring within each time bin. Bottom: CCH z-scored relative to shuffled vSWR onset times. Z-score 0 reflects mean of shuffles. Error bars indicate s.e.m. **f**, Mean CCH showing high synchrony of awake SWRs between pairs of tetrodes within dCA1 (top, n=5 rats) or vCA1 (bottom, n=3 rats with >1 tetrode in vCA1), z-scored relative to shuffled event times. SWRs here were detected on individual tetrodes. Error bars indicate s.e.m. **g**, Change in mean dSWR rate on rewarded vs. error trials (well visits), calculated per time spent immobile at reward wells (minimum 1 s). Each point represents the mean SWR rate across animals ± s.e.m. within that learning stage (n=4 rats in Acquisition stages 0.7-0.8, >0.8; n=5 rats all other stages; see Online Methods), defined by each animal’s probability of correct performance on each trial (see Fig. 1c, Extended Data Fig. 2). Learning stage is used here as a proxy for novelty; the task and environment are both novel when Acquisition performance of the first sequence is <0.3. For Switch performance, SWR rate is shown only for the second sequence when it was rewarded. Learning stage means are fit by an exponential decay function for both rewarded and error trials (dotted lines indicate 95% confidence bounds on the fit). **h**, Similar to **g**, but for vSWR rate in each learning stage (n=4 rats in Acquisition stages 0.7-0.8, >0.8; n=5 rats all other stages).

We examined awake SWRs detected in dH and vH during periods of immobility on the task, which occurred primarily at the reward wells (example in Fig. 1d). Both dH and vH SWRs (dSWRs and vSWRs) showed the expected spectral properties and increases in multiunit activity^16, 20^, indicating strong, transient activation of local hippocampal networks at the times of SWRs (Extended Data Fig. 3).

Strikingly, despite the existence of dorsoventral connectivity within the hippocampus^21^ and observations of occasional synchrony between dSWRs and vSWRs during sleep^20^, dSWRs and vSWRs occurred asynchronously during awake immobility on the maze (Fig. 1d,e). Only ∼3.7% of dSWRs occurred within 50 ms of a vSWR, which was no more than expected from a shuffle of SWR times (Fig. 1e, Extended Data Fig. 4a,c). This asynchrony was in stark contrast to the prominent synchrony between pairs of recording sites within dH or within vH (Fig. 1f, Extended Data Fig. 4b), indicating temporally separable dH and vH outputs to downstream brain areas during SWRs.

DSWRs and vSWRs also showed very different patterns of modulation by novelty and reward. As expected given previous findings^15–17, 19^, dSWRs were more prevalent when the environment was novel and following receipt of reward (Fig. 1g, Extended Data Fig. 4d). Given the strong anatomical projections from the vH to limbic brain areas involved in reward processing^5, 6, 8–11^, we expected that vSWRs would show a similar or greater level of reward modulation. Instead, vSWRs maintained a similar rate on rewarded and error trials and were not enhanced during early novelty (Fig. 1h, Extended Data Fig. 4d). The onset time of dSWRs and vSWRs also differed: while dSWRs began only after receipt of reward following initial learning, vSWRs were detected as soon as the animal stopped moving (Extended Data Fig. 4e,f). Thus, dSWRs and vSWRs are differently regulated by novelty and reward.

The temporal separation of awake dSWRs and vSWRs provided the opportunity to determine whether these events differentially engaged the NAc. Previous work reported activation of NAc neurons during dSWRs in sleep^3, 4^. It remains unknown whether NAc neurons are engaged during awake dSWRs or during either awake or sleep vSWRs. Importantly, it is also unknown whether dSWRs and vSWRs engage similar or contrasting NAc populations. To sample the respective target regions of the sparse dH projection and the much more prominent vH projection^6–10, 12^, we recorded from both the NAc core and medial shell (Extended Data Fig. 1d). We then classified NAc single units into putative medium spiny neurons (MSNs) and fast-spiking interneurons (FSIs) (Extended Data Fig. 5a) and examined their activity around the times of awake dSWRs and vSWRs.

We found that 51% of MSNs either significantly increased or decreased their firing rates around the times of dSWRs and/or vSWRs. Strikingly, the observed firing rate changes were often opposite for dSWRs and vSWRs, such that 10.6% of cells were significantly dSWR-activated and vSWR-suppressed (D+V-) or dSWR-suppressed and vSWR-activated (D-V+) (Fig. 2a,b). Crucially, this fraction was significantly larger than would be expected by total chance overlap of the D+/V- and D-/V+ subgroups, while the fraction of co-positively modulated cells (3.2% D+V+) was not greater than chance. In addition, many cells were significantly modulated during only dSWRs or vSWRs but not both (Fig. 2b). Across the full population of MSNs (Fig. 2c, see also Extended Data Fig. 6a), we found a significant anti-correlation in SWR modulation amplitudes (Fig. 2d), demonstrating that dSWRs and vSWRs are consistently associated with opposite activity changes at the level of individual NAc MSNs. We verified that these results held when the small number of coincident dSWRs and vSWRs were excluded (Extended Data Fig. 6e-g) and when we applied a conservative criterion (Extended Data Fig. 5c-j) to ensure that each cell was included only once (Extended Data Fig. 6h-j). We also noted that MSN activity changes predominantly followed dH or vH neuronal activation during SWRs, suggesting hippocampus to NAc information flow (Extended Data Fig. 6b).

**Figure 2.**
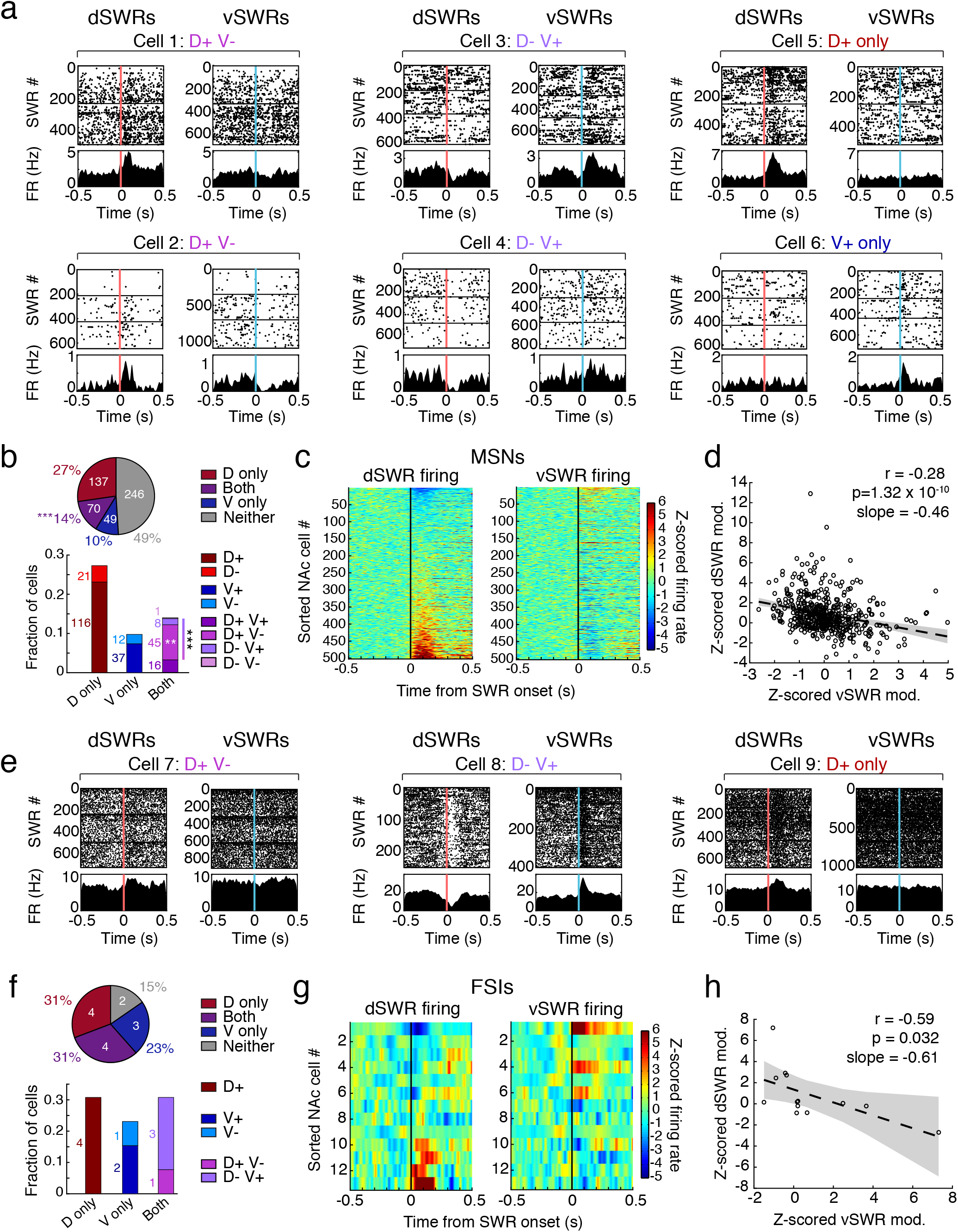
Opposing patterns of NAc modulation during awake dH vs. vH SWRs. **a**, Examples of single NAc MSNs showing significant modulation during awake SWRs on the Multiple-W task. Spike rasters and peri-event time histograms (PETHs) are aligned to the onset of dSWRs (left within each cell, pink line) or vSWRs (right within each cell, blue line). Horizontal lines separate maze epochs. Categories at the top of each cell indicate directions of significant modulation (positive or negative). All modulations in these examples are significant at p<0.01 (shuffle test). **b**, Proportions of significantly SWR-modulated NAc MSNs. Top: fractions modulated during dSWRs only (D only), vSWRs only (V only), Both, or neither dSWRs nor vSWRs (Neither) regardless of the direction of modulation. Cell counts are shown in white. Significantly more cells are modulated during Both than would be expected by chance overlap of dSWR- and vSWR-modulated cells (***p=5.44×10^4^, z-test for proportions). Bottom: directional modulation of MSNs. Cell counts are shown next to each bar. The fractions of D+V-cells alone and total “opposing” cells (gradient bar, D+V- and D-V+) are higher than would be expected by chance (**p=0.0017 and ***p=6.12×10^4^, respectively, z-tests for proportions). All fractions are out of 502 MSNs that fired at least 50 spikes around both dSWRs and vSWRs, from all 5 rats. **c**, NAc MSN population shows opposing modulation during dSWRs vs. vSWRs. Left: dSWR-aligned z-scored PETHs for each MSN ordered by its modulation amplitude (mean z-scored firing rate in the 200 ms following SWR onset). Right: vSWR-aligned z-scored PETHs for the same ordered MSNs shown on the left. Z-scores are calculated within cell relative to the pre-SWR period (−500 to 0 ms). **d**, Anti-correlation between dSWR and vSWR modulation amplitudes of MSNs. Each point represents a single cell. Dotted line and shaded regions represent a linear fit with 95% confidence intervals. Pearson’s correlation coefficient (r), p-value and slope are shown in upper right. **e**, Examples of single NAc FSIs showing significant modulation during SWRs. Format and modulation categories as in **a**, top row. All modulations are significant at p<0.01 (shuffle test). **f**, Proportions of significantly SWR-modulated FSIs after exclusion of any potential duplicates from holding the same cell across days, which could bias this small population of cells (see Online Methods). Fractions are out of 13 FSIs from 5 rats. Similar to **b**. **g**, NAc FSI population shows opposing modulation during dSWRs vs. vSWRs. Similar to **c**. **h**, Anti-correlation between dSWR and vSWR modulation amplitudes of FSIs. Similar to **d**.

The same opposing patterns of modulation were seen when we examined FSIs, applying the same criterion to ensure that each cell was included only once (Extended Data Fig. 5c-j). The majority of FSIs were SWR-modulated (85%), and we identified FSIs that were D+V-, D-V+, or only D+ or V+, but none that were D+V+ (Fig. 2e,f). The FSI population as a whole showed anti-correlated modulation during dSWRs versus vSWRs (Fig. 2g,h, Extended Data Fig. 6c,d). As individual FSIs innervate large populations of MSNs^22^, FSIs may play an important role in coordinating NAc responses to hippocampal inputs^7^. We also found that SWR-modulation of both MSNs and FSIs was anatomically distributed in a pattern consistent with reported dH and vH projections to the NAc^6–10, 12^, with V+ neurons present in both the medial shell and parts of the core and D+ neurons restricted to the core (Extended Data Fig. 7).

The Multiple-W task involves reward associations with particular spatial paths, allowing us to test the prevailing hypothesis that vH would most strongly engage valence-related representations downstream^10, 11, 23–26^, including reward associations in the NAc. We speculated that dSWR- and vSWR-activated NAc representations may be distinctly related to the task, since when one is activated the other is typically suppressed.

Consistent with the possibility of distinct representations, we identified multiple differences between dSWR- and vSWR-activated NAc firing patterns outside of SWRs. Surprisingly, these differences highlighted a specificity for spatial-reward information in the dSWR-activated neurons. To examine the dSWR- and vSWR-activated populations independently, we grouped together all cells that were D+ (only D+ or D+V-) and separately, all cells that were V+ (only V+ or D-V+), excluding the D+V+ cells. We then examined the firing patterns of each population on the six rewarded trajectories across the two alternation sequences. Each trajectory included both time spent at the reward well starting the trajectory (“well,” excluding spikes during SWRs) and time spent during movement between reward wells (“path”; Fig. 3a).

**Figure 3.**
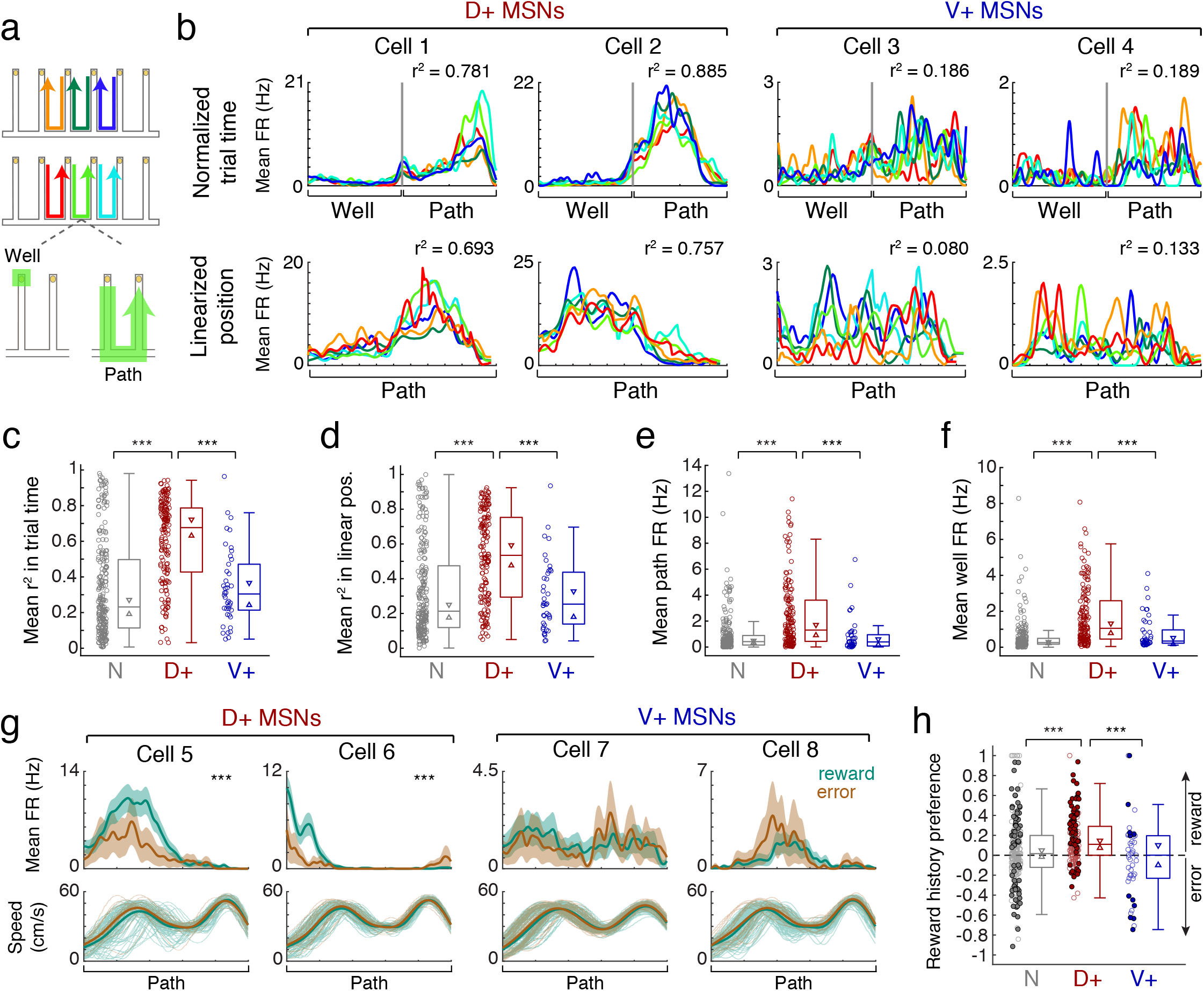
Selective encoding of task-related information in the dH-NAc network. **a**, Schematic of the 6 rewarded trajectories across Sequence A and B, defined spatially by start and end reward well and directionally by left (top) or right (middle) movement between wells. Bottom: An example trajectory split into its well and path components, where well is the time from nosepoke to turnaround (departure) and path is the time from turnaround to next nosepoke. **b**, Example NAc MSNs from the D+ and V+ populations showing high (D+, left) vs. low (V+, right) firing similarity (r^2^) across trajectories. Trajectories are color coded according to the key in **a**. Cell numbers do not correspond to Fig. 2. Top row: trajectories are plotted as a function of normalized trial time. Grey vertical line marks 100% of the well period and beginning of the path period, and the end of the path period starts a new well visit. Bottom row: trajectories are plotted as a function of linearized position (one-dimensional distance in cm) from each trajectory’s start well to end well. Note that well times fall into the first and last position bins of the path since there is no position change at the well. X-axis for each linearized position plot covers ∼220 cm. Pearson’s r^2^ across all possible pairs of trajectories is shown in upper right. Controls for behavioral variability were included (see Online Methods). **c, d**, Distributions of mean r^2^ in normalized trial time (**c**) and linearized position (**d**) by SWR-modulation category. **c**, N (n=226 cells) vs. D+ (n=154 cells): ***p=5.45×10^19^; D+ vs. V+ (n=42 cells): ***p=6.33×10^9^. **d**, N (n=222 cells) vs. D+ (n=152 cells): ***p=3.17×10^12^; D+ vs V+ (n=40 cells): ***p=5.46×10^7^ (Wilcoxon rank-sum tests). Circles are individual cells, boxes show interquartile range, horizontal lines mark the median, triangles mark the 95% confidence interval of the median, whiskers mark non-outlier extremes. **e, f**, Distributions of mean firing rate outside of SWRs during maze epochs, during path (**e**) and well periods (**f**), by SWR-modulation category. **e**, N (n=226 cells) vs. D+ (n=154 cells): ***p=1.57×10^12^; D+ vs. V+ (n=42 cells): ***p=5.70×10^6^. **f**, N vs. D+: ***p=4.94×10^22^; D+ vs. V+: ***p=3.69×10^5^; V+ vs. N: p=0.024, n.s. with Bonferroni correction for multiple comparisons (Wilcoxon rank-sum tests). Circles and boxes as in **c,d**. **g**, Example NAc MSNs from the D+ (left) and V+ (right) populations showing path firing patterns as a function of reward history, defined as whether or not the previous well visit was rewarded. Top: firing rate of each cell (mean ± s.e.m.) on all paths following a reward (teal) vs. all paths following an error (brown). Reward history preference (see Online Methods): Cell 5: 0.30, Cell 6: 0.47, Cell 7: 0.025, Cell 8: −0.19. ***p<0.001 indicates cells with significantly higher firing rate on paths following reward compared to error (one-tailed permutation test). Bottom: faded lines indicate speed profiles of individual paths following reward and error, thick lines indicate mean speeds. **h**, Distributions of reward history preference by SWR-modulation category. Filled circles indicate significantly reward-preferring (>0) or error-preferring (<0) cells, open circles indicate non-significant cells. The D+ population (n=160 cells) is significantly shifted toward positive values (p=8.18×10^13^, one-tailed signed-rank test) in additional to being significantly more reward-preferring than the N (n=241 cells, ***p=2.63×10^5^) and V+ (n=45 cells, ***p=0.00041) populations.

First, we found that D+ MSNs tended to fire very similarly across distinct trajectories as a function of both normalized trial time and linearized position (see Online Methods). We quantified firing similarity as a mean coefficient of determination (r^2^) across pairs of trajectories. D+ MSNs with high r^2^ values were “tuned” to the same relative point of progression through each trajectory in both time and distance, regardless of actual spatial location or movement direction (examples in Fig. 3b, left). This preferred trajectory stage varied across D+ cells, such that D+ population activity spanned the full extent of each trajectory (Extended Data Fig. 8a). Such patterns have been reported in dorsal and ventral striatum^27–29^ and are consistent with dH input^30^, but they have not been previously linked to dSWR activation. Moreover, these D+ MSN firing patterns mirror the subset of medial prefrontal cortical neurons that are activated during dSWRs^31^, consistent with a role for dSWRs in broadly engaging task-related representations that generalize across different spatial paths.

By contrast, many V+ MSNs showed sparser firing patterns that were largely uncorrelated between distinct trajectories (Fig. 3b, right) and even between traversals on the same trajectory (Extended Data Fig. 9a,b). Across the population, D+ MSNs showed significantly higher firing similarity across trajectories compared to both the V+ MSNs and unmodulated (N) MSNs (Fig. 3c,d). Furthermore, the D+ population had much higher mean firing rates on both the path and well components suggesting greater task engagement (Fig. 3e,f, Extended Data Fig. 8b-e). Increased correlations across trajectories persisted when we controlled for firing rates (Extended Data Fig. 8f,g). These findings indicate that, in the context of our task, D+ cells (and D+V+ cells, Extended Data Fig. 8h) are much more active and express clear task-related firing properties that are not evident in cells that are V+ and not D+ (see also Extended Data Fig. 9c-g).

We also observed a strong and differential effect of reward history on D+ MSNs, consistent with a role for these neurons in representing the outcome of the previous choice^8, 9^. We computed a reward history preference for each MSN from the difference in its mean firing rate curve on paths following a rewarded well visit versus an unrewarded visit (error), controlling for movement speed and the influence of upcoming choice (Fig. 3g,h; Extended Data Fig. 8i-l). We found that the D+ population fired more on paths following reward than following an error, demonstrating a clear reward history preference that was not seen in the V+ or unmodulated populations (Fig. 3h). At the same time, we were surprised to find an overall lack of enhanced firing during receipt of reward at the reward sites, given previous work suggesting reward-site-specificity for NAc neurons activated during dSWRs in sleep^3, 4^. While we found individual examples of MSNs that had significantly higher firing rates during rewarded as opposed to unrewarded times at the wells, we also found cells that showed the opposite pattern, and neither the D+ nor V+ populations were enriched for cells showing reward-specific firing at the wells (Extended Data Fig. 9h-k). These results suggest that a learned relationship between spatial paths and reward outcomes is selectively reflected in D+ MSNs.

Finally, we asked whether D+ and V+ neurons constitute separate networks across both waking task performance and sleep, as we would expect if these physiological subtypes reflect distinct anatomical networks. During sleep there is greater synchrony between dSWRs and vSWRs^20^ (∼6.7% of dSWRs, Extended Data Fig. 10), so we first excluded all pairs of dSWRs and vSWRs that occurred within 250 ms of each other. In the remaining, temporally isolated SWRs in sleep, we found that the proportion of single MSNs showing opposing modulation was again greater than chance (Fig. 4a). Furthermore, the population-level opposition during dSWRs versus vSWRs remained apparent, particularly for MSNs (Fig. 4b-f).

**Figure 4.**
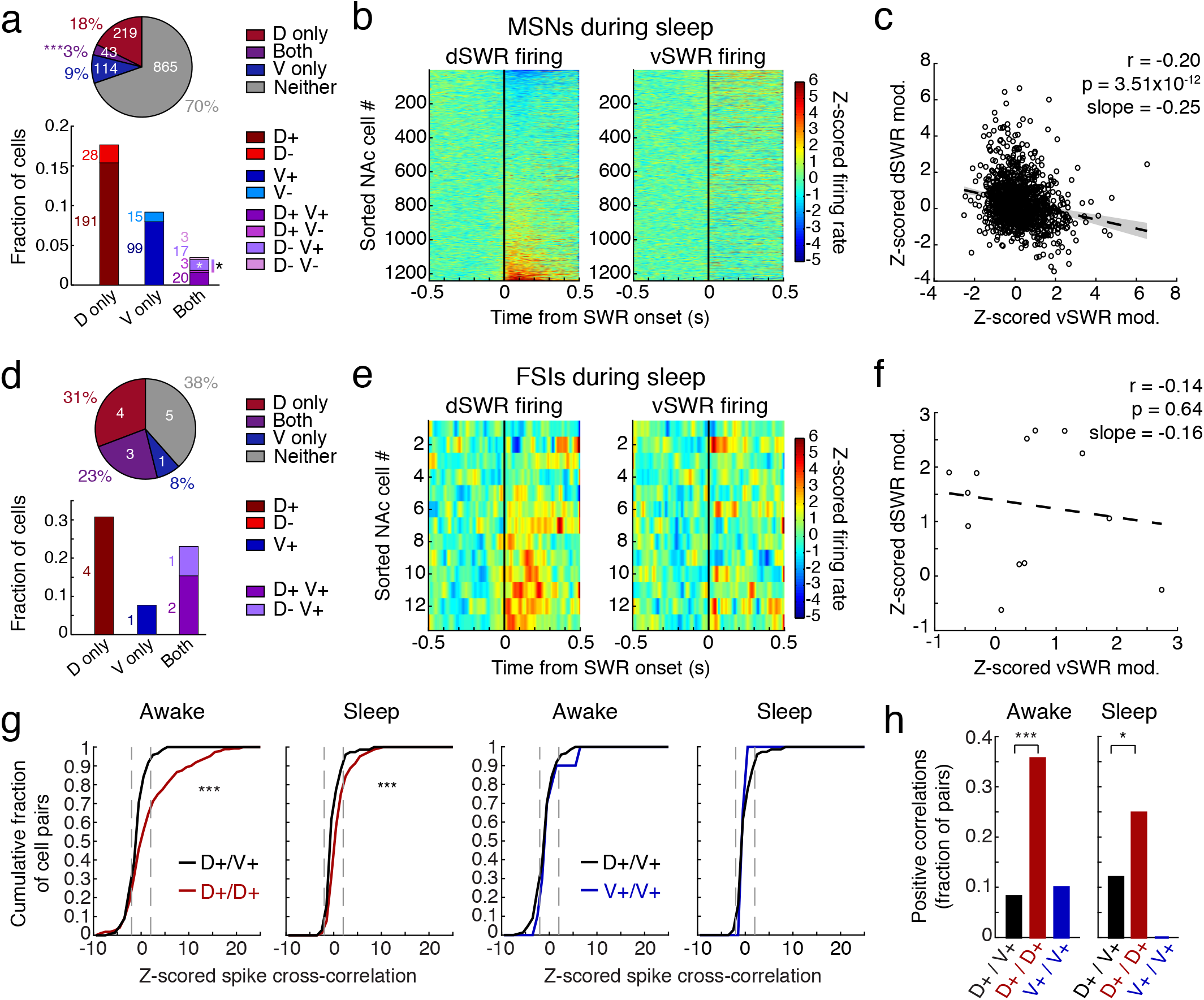
Hippocampal-NAc network patterns are maintained during sleep. **a**, Proportions of NAc MSNs showing significant modulation during asynchronous (more than 250 ms apart) dSWRs and vSWRs in sleep, similar to Fig. 2b. Top: fractions of modulated MSNs regardless of the direction of modulation. Cell counts are shown in white. Significantly more cells are modulated during Both than would be expected by chance overlap of dSWR- and vSWR-modulated cells (***p=1.50×10^4^, z-test for proportions). Bottom: directional modulation of NAc MSNs. Cell counts are shown next to each bar. The fraction of D-V+ cells alone and total “opposing” cells (gradient bar, D+V- and D-V+) are higher than would be expected by chance (*p=0.012 and *p=0.037, respectively, z-tests for proportions). All fractions are out of 1241 MSNs that fired at least 50 spikes around both dSWRs and vSWRs in sleep, from all 5 rats. **b**, NAc MSN population activity shows opposing modulation during asynchronous dSWRs and vSWRs in sleep, similar to Fig. 2c. Left: dSWR-aligned z-scored PETHs for each MSN ordered by its modulation amplitude. Right: vSWR-aligned z-scored PETHs for the same ordered MSNs shown on the left. **c**, Anti-correlation between dSWR and vSWR modulation amplitudes of MSNs, similar to Fig. 2d. Each point represents a single cell. Dotted line and shaded regions represent a linear fit with 95% confidence intervals. Pearson’s correlation coefficient (r), p-value and slope are shown in upper right. **d,** Similar to **a**, but for fractions of FSIs showing significant modulation during asynchronous SWRs in sleep, after removal of potential duplicate cells. Fractions are out of 13 FSIs. **e,** Similar to **b**, but for FSI population in sleep. **f,** Similar to **c**, but for FSI population in sleep. **g,** Cumulative distributions of z-scored spike cross-correlations (mean at zero lag ±10 ms) between pairs of MSNs outside SWRs. Left two plots: pairs of D+ MSNs (D+/D+) vs. pairs of D+ and V+ MSNs (D+/V+) during awake movement (far left) and during sleep (second from left). The D+/D+ distribution is significantly shifted to the right of the D+/V+ distribution in both states (awake ***p=5.01×10^7^; sleep ***p=2.86×10^5^; one-tailed Wilcoxon rank-sum tests). Right two plots: pairs of V+ MSNs (V+/V+) vs. D+/V+ pairs, during awake movement (second from right) and during sleep (far right). The V+/V+ distribution is not significantly different from the D+/V+ distribution in either state. D+/V+ distributions are repeated from left to right for clarity. D+ and V+ categories are assigned based on awake SWR modulation, excluding cells that are both D+V+. Sleep cross-correlations were only calculated for awake-SWR-modulated cells also active in sleep (awake n=272 D+/D+ pairs, 146 D+/V+ pairs, 10 V+/V+ pairs; sleep n=161 D+/D+ pairs, 75 D+/V+ pairs, 3 V+/V+ pairs). **h**, Fraction of cell pairs exhibiting positive spike cross-correlations (≥2 z-scores, mean at zero lag ±10 ms). Awake: D+/D+ vs. D+/V+: ***p=1.11×10^9^; D+/D+ vs. V+/V+: p=0.094. Sleep: D+/D+ vs. D+/V+: *p=0.025; D+/D+ vs. V+/V+: p=0.32 (z-tests for proportions).

Patterns of co-firing outside of SWRs provided further evidence for distinct networks. During both awake behavior and sleep, pairs of D+ MSNs showed a stronger tendency to be coactive than D+/V+ pairs. No such difference was seen for V+ MSN pairs as compared to D+/V+ pairs (Fig. 4g,h), perhaps because of the overall low levels of activity of V+ neurons in our task (Fig. 3e,f). These findings indicate that the D+ MSN population constitutes a specific network distinct from the V+ MSN population.

Our findings identify two distinct and opposing networks in the NAc, one activated during dSWRs and another activated during vSWRs. These two networks are activated at different times, and only the dSWR-activated (D+) network showed strong encoding of information related to spatial experience and past reward, consistent with a recent report^7^. The V+ NAc neurons, in contrast, were not evidently tuned to specific features in our appetitive task. As vH has been shown to encode both broad spatial context^23, 32, 33^ and aversive experiences^24, 26^, neither of which was relevant for our task, we suggest that the vH-NAc network could be specialized for associations not engaged in our task such as those between overall context and valence. While the specific role of the vH-NAc network remains unclear, our findings establish an opposition between the dH-NAc and vH-NAc networks that is well suited to support the processing of different aspects of experience at different times.

## Author contributions

M.S. and L.M.F. designed the study. M.S. and H.R.J. conducted experiments. M.S. analyzed the data. M.S. and L.M.F. wrote the paper with input from H.R.J.

## Acknowledgements

We thank J. Berke, H. Fields, M. Kheirbek, M. Brainard, A. Gillespie, A. Joshi, A. Comrie, M. Coulter, J. Yu, and K. Kay for comments on the manuscript, and members of the Frank laboratory for useful discussions. We additionally thank K. Kay for contributing key analysis code, J. Chung and J. Magland for development of drift tracking in Mountainsort, and I. Grossrubatscher, V. Kharazia, and E. Miller for technical assistance. This work was supported by Howard Hughes Medical Institute (L.M.F.), Simons Foundation Collaboration for the Global Brain Grants 521921 and 542981 (L.M.F.), National Institute of Mental Health Ruth L. Kirschstein National Research Service Awards F31MH111214 (M.S.) and F30MH115582 (H.R.J.), and National Institute of General Medical Sciences Medical Scientist Training Program Grant T32GM007618 (H.R.J.). The content is solely the responsibility of the authors and does not necessarily represent the official views of the National Institutes of Health.

## Methods

### Animals, implants, and behavior

All procedures were in accordance with guidelines from the University of California San Francisco Institutional Animal Care and Use Committee and US National Institutes of Health. Five male Long-Evans rats (500-650 g, 5-8 months old) were food restricted to 85% of their baseline weight and pre-trained to run back and forth on a 1 m long linear track for liquid reward (evaporated milk plus 5% sucrose), delivered automatically from reward wells at the ends of the track. Animals were incrementally introduced to a delay between well entry (nosepoke) and reward delivery of up to 2 seconds. After animals learned to alternate consistently for at least ∼30 well visits per 5 min (4-6 days), they were switched to an *ad libitum* diet and then surgically implanted with microdrive arrays.

Each microdrive housed a maximum of 28 independently movable tetrodes in a custom 3D-printed drive body (PolyJetHD Blue, Stratasys Ltd.) cemented to 3 stainless steel cannulae at fixed relative positions, targeting NAc vertically (8-12 tetrodes) and dH (6-7 tetrodes) and vH (9-13 tetrodes) at a 12° angle from vertical (tilted mediolaterally). NAc and dH tetrodes were made of 12.7 µm-diameter nichrome (Sandvik), while vH tetrodes were made of 12.7 µm nichrome, 12.7 µm tungsten, or 20 µm tungsten (California Fine Wire). Tetrode ends were plated with gold to a final impedance of ∼240-330 kOhms. The microdrive was stereotaxically implanted over the right hemisphere such that the center of each cannula was targeted to the following coordinates relative to the animal’s Bregma: dH: AP −3.9-4.0 mm, ML +1.7 mm; vH: AP −5.6-5.7 mm, ML +4.0 mm; NAc: AP +1.3-1.4 mm, ML +1.3 mm (Rat 1 vH: AP −5.75, ML +4.1, oval-shaped cannula). The approximate AP/ML spread of tetrodes in each area was defined by the inner radius of each cannula as follows: dH: ±0.49 mm, vH: ±0.87 mm, NAc: ±0.60 mm. A ground screw was inserted in the skull above the right cerebellum as a global reference.

While animals recovered from surgery, tetrodes were manually adjusted over ∼2-3 weeks to their target depths relative to brain surface (dCA1: ∼2.2-3.3 mm, 12° angle; vCA1: ∼7.0-8.3 mm, 12° angle; NAc: ∼5.4-7.5 mm, 0° angle), using electrophysiological landmarks such as unit density and SWR amplitude. Each rat was then food restricted again and re-trained on the linear track for 4-6 days with neural recording (not analyzed in this study). Animals were then introduced to the Multiple-W task (Fig. 1b), a version of which has been described previously^19^. Tetrodes were sometimes advanced a small amount after the conclusion of the day’s recording on a case-by-case basis to improve cell yield.

The Multiple-W track consisted of six 76 cm arms spaced ∼36 cm apart at their midpoints, with 3 cm high walls, connected to a “back” which extended past the first and sixth arm by 14 cm on each side (to mimic the availability of a right and left turn from these arms), and elevated 76 cm off the ground. On each day, the animal experienced three 20 min “run” (maze) epochs on the track flanked by four 20-45 min sleep epochs in a separate high-walled rest box; only in rare cases (2 epochs each for Rats 1-2, 1 epoch each for Rats 3-4) were there four run epochs. The track was separated from the experimenter by an opaque black curtain, and the white walls of the room were marked with black distal cues of various shapes. Each arm contained a visually identical reward well connected to milk tubing, and milk was run through each well at the beginning of the day to create similar olfactory cues in all wells.

In each run epoch, the animal was placed at the back of the center arm of the rewarded sequence and was required by trial-and-error to find the 3 rewarded wells and figure out the alternation sequence between them, Sequence A (SA) or Sequence B (SB). Trials are defined as well visits. A visit to the center well of the sequence (well 3 in SA, well 4 in SB) was rewarded if the animal came from any other well. If a center visit was the first of the epoch or followed an error to a non-sequence arm, the animal could initiate an “outer” well visit to either of the center-adjacent wells to get reward. If a center visit followed a visit to a center-adjacent well, the animal had to then visit the opposite center-adjacent well, requiring hippocampal-dependent memory of the previous trial^34^. For example, a correct series of trials for SA would be 3-2-3-4-3-2; for SB, 2-1-2-3-2-1. Consecutive visits to the same well were counted as errors, such that chance performance was defined as 1 out of 6 (0.167). The nosepoke at each well was detected by an infrared beam break, which automatically triggered liquid reward delivery (105 µL evaporated milk plus 5% sucrose) via a syringe pump (OEM Systems) after a 2 second delay, and the animal’s departure from the well was self-paced.

During the “Acquisition” phase (5-9 days), the same sequence was rewarded on every epoch: 3 animals acquired SA and 2 animals acquired SB. When the animal achieved greater than 80% correct performance on the Acquisition sequence for at least 1 epoch (assessed in real time as an epoch average), the novel sequence was introduced in the second epoch of the first “Switch” day. Only Rat 1 failed to reach 80% correct but was advanced to the Switch phase after achieving >75% correct and one full week of training as in ref. 19; this rat was thus excluded from the 70-80% and >80% Acquisition performance bins in Fig. 1g,h. In the Switch phase, the rewarded sequence was switched on each run epoch, such that the starting sequence of each Switch day was also alternated (8-10 days). The goal was to continually promote adaptive, spatially-guided reward-seeking behavior, and the overlap between arms of the two sequences allowed us to compare rewarded and error trials on the same spatial paths.

### Data collection and processing

Spiking, local field potential (LFP), position video, and reward well digital inputs and outputs were collected using the NSpike data acquisition system (L.M.F and J. MacArthur, Harvard Instrumentation Design Laboratory). For Rats 1-3, LFP data were collected at 1500 Hz sampling rate and digitally filtered online at 1-400 Hz (2-pole Bessel for high- and low-pass). Spikes were sampled at 30 kHz and saved as snippets of each waveform, filtered at 600-6000 Hz for hippocampus and 600-6000 Hz (2 rats) or 300-6000 Hz (1 rat) for NAc. For Rats 4-5, LFP and spikes were collected continuously at 30 kHz and filtered online at 1-6000 Hz, with post-hoc filtering applied in Matlab to extract LFP and spike waveforms using the same parameters as above (300-6000 Hz for NAc spikes). Subsets of spike data were collected as snippets in these animals to verify our post-hoc filtering. Note that for LFP and spike waveforms, negative voltages are displayed upward. All LFP and spikes were collected relative to local references lacking spiking activity, which were themselves referenced to cerebellar ground: for dH tetrodes, the reference tetrode was located in corpus callosum (4 rats) or deep cortex with no units (1 rat); for vH, in ventral corpus callosum (1 rat) or in white matter at the ventrolateral edge of the midbrain (internal capsule or optic tract; 4 rats); for NAc, typically in corpus callosum, the lateral ventricle, or anterior commissure. Overhead video of the track, collected at 30 frames/s, allowed us to track the animal’s position via an array of infrared diodes attached to the top of the headstage, a few cm above the rat’s head.

Spike sorting was performed using a combination of manual clustering in Matclust (Mattias Karlsson; Rats 1-3) and automated sorting with manual curation in Mountainsort (Flatiron Institute; all data for Rats 4-5, individual days for Rats 1-3). Cells were clustered within epochs but tracked across all run and sleep epochs for which they could be isolated; with Mountainsort, this was done with a drift-tracking extension of the core pipeline and manual merging as needed^35^. In Matclust, clustering was performed in amplitude and principal component space, and only well-separated units with clear refractory periods in their ISI distributions were accepted. In Mountainsort, we generally accepted clusters with isolation score >0.96, noise overlap <0.03 (median isolation score ∼0.995, median noise overlap ∼0.002), and clear separation from other clusters in amplitude and principal component space. The similarity of cluster quality between Mountainsort and Matclust was verified manually on a subset of the data and has been extensively verified in previous work^36^. The same pattern of SWR-modulation of NAc cells was observed within each animal (data not shown), indicating that our results were not due to unit clustering in certain animals.

### Histology

At the conclusion of the experiment, animals were anesthetized with isoflurane and small electrolytic lesions were made at the end of each tetrode to mark recording locations (30 µA of positive current for 3 seconds, on 2-4 channels of the tetrode). The animal recovered for 24 hours to allow gliosis, and was then euthanized with pentobarbitol and perfused transcardially with PBS followed by 4% paraformaldehyde in 0.1M PB. The brain was post-fixed in 4% paraformaldehyde, 0.1M PB *in situ* for at least 24 hours, followed by removal of the tetrodes and cryoprotection in 30% sucrose in PBS. Brains were embedded in OCT compound and sectioned coronally at 50 µm thickness. Tissue was either Nissl stained using cresyl violet, or for a subset of dH sections, immunostained for RGS14, a marker of CA2, using previously described methods^37^.

To reconstruct recording sites (Extended Data Fig. 1), evenly spaced plates from the Paxinos and Watson Rat Atlas (2007), which is based on Wistar rats, were stretched and modified to align to representative sections from each brain region, using landmarks such as the ventricles, corpus callosum, and hippocampal pyramidal layers as guides. These modified plates were then treated as atlases to align the remaining sections and recording sites across animals.

### Data analysis

All analyses were performed using custom code written in Matlab.

### Behavioral analysis

The animals’ task performance was analyzed using a state space algorithm^38^ which estimates the probability that the animal is performing accurately according to Sequence A or B on each trial. This algorithm provides 90% confidence intervals which reveal when the animal is performing one sequence significantly better than the other. All trials in the Acquisition phase were analyzed together and background probability was set at chance (0.167), so that behavior of all animals could be compared from a similar starting point. During the Switch phase, each epoch was estimated independently with an unspecified background probability to get the most accurate representation of the animals’ behavior; this means that occasionally the behavioral state could “jump” at an epoch boundary. The mode of the probability distribution was used to assign trials into performance stages for the SWR rate analysis in Fig. 1g,h, which yielded a different number of trials per stage from each animal.

### SWR detection

SWRs were detected in dCA1 and vCA1 in 4 rats (Rats 2-5), and dCA1 and ventral CA3 in 1 rat (Rat 1), using methods described previously^37^. Briefly, each tetrode’s LFP was filtered to the ripple band at 150-250 Hz, the ripple amplitude was squared, summed across tetrodes (3 per animal in dH, only 1 per animal in vH, as this was the minimum number present in all animals), and smoothed with a 4 ms s.d., 32 ms wide Gaussian kernel. We then took the square root of this trace as the power envelope to detect excursions greater than 2 s.d. of the mean power within an epoch, lasting at least 15 ms. Tetrodes were chosen based on ripple band power and proximity to the center of the pyramidal layer. The SWR start time (when the envelope first crosses the threshold) was used as the event detection time. For spiking, characterization, and cross-correlation analyses, we excluded SWRs that occurred within 0.5 s of a previous SWR (i.e. chained SWRs). SWRs were only included for all analyses if detected at head speeds <4 cm/s.

As a control, we also detected “noise ripples,” events in the 150-250 Hz band that exceeded a 2 s.d. threshold on our reference tetrodes for dH and vH (which were not in the hippocampus). These events are highly unlikely to be SWRs, but instead may reflect muscle artifacts or other high-frequency noise. For all analyses of SWRs other than NAc spiking modulation (to include the maximum number of NAc spikes), we excluded SWRs with start times occurring within 100 ms of a “noise ripple” on the local reference.

### Behavioral state definitions

During run epochs, periods of “immobility” were defined as times with a head movement speed <4 cm/s calculated as the derivative of the smoothed position data from the headstage-mounted diodes. We defined “sleep” as periods of immobility in sleep epochs that occurred >60 s after any movement at >4 cm/s. To calculate overall sleep SWR rate in Extended Data Fig. 3, NREM sleep periods were defined by exclusion of REM sleep as defined previously^37^. Specifically, REM periods were detected as times when the ratio of Hilbert amplitudes of theta (5–11 Hz) to delta (1–4 Hz), referenced to cerebellar ground, exceeded a per-animal threshold of 1.4-1.7 for at least 10 s.

### Characterization of SWR properties

To characterize the spectral properties of dSWRs and vSWRs, multi-tapered spectrograms of the raw LFP triggered on SWR start times were generated using the Chronux toolbox (mtspecgramtrigc, sliding 100 ms window with 10 ms overlap, bandwidth 2-300 Hz), and z-scored to the mean power in each epoch before averaging across epochs and days. To approximate the peak ripple frequency, a slice of this spectrogram was taken at the time of peak ripple power per animal. For the remaining properties described in Extended Data Fig. 3: we defined SWR amplitude as the minimum threshold in s.d. that would be required to detect the event (see above). SWR duration is the time between first threshold crossing and return of the envelope to the mean. Mean epoch SWR rate was calculated for all immobility periods in run epochs and all NREM sleep periods during sleep epochs.

### SWR cross-correlation

Cross-correlations between vSWRs and dSWRs were performed within day, using dSWRs as the reference, in 50 ms bins up to 0.5 s lag, and were normalized to the number of dSWRs in each day before averaging across days and animals. To create a z-scored version of the cross-correlation histogram, vSWR event times were circularly shuffled 1000 times within immobility periods (by a random amount up to ±half the mean immobility period length) to create 1000 shuffled histograms. The real cross-correlation values were z-scored relative to the distribution of shuffled values within each bin, such that a z-score of 0 indicates that the real data is no different than the mean of the shuffles.

### SWR rate relative to reward and novelty

In Fig. 1g,h, we calculated the rate of SWR events per time spent immobile after reward delivery time on individual rewarded or error trials, and then averaged those rates across trials in each learning stage from each animal. Trials with less than 1 s spent immobile were excluded. In Extended Data Fig. 4d, this rate was averaged across trials within each run epoch for each animal. In Extended Data Fig. 4e,f, we calculated SWR rate in 200 ms bins from 0 to 5 s after nosepoke. We subsampled rewarded and error trials based on speed by excluding any trial where the animal spent more than 5 position samples (150 ms) moving faster than 4 cm/s (allowing for some jitter of head position), from 1.5 s after nosepoke to the end of the 5 s window. As the animal’s retreat from the reward well is self-paced, this greatly reduced the number of included error trials and focused exclusively on error trials when the animal waited at the well beyond the expected reward delivery time. We also excluded any bins that were not below the speed threshold, as SWRs could not be detected in these bins according to our criteria. SWR rate per bin was then calculated per the number of included bins in each animal, and we required at least 2 s total of data per bin (10 accepted bins) to calculate a rate across animals.

### Unit inclusion

Only NAc units firing at least 100 spikes in a given epoch were included in the current study (865 total NAc units in run, 1678 units in sleep, from all five rats across days). Additional inclusion criteria were applied per analysis.

### Putative cell type classification

NAc single units were classified similar to methods described previously^39^, using mean firing rate, mean waveform peak width at half-maximum, mean waveform trough width at half-minimum, and ISI distribution. These values were averaged across epochs when a cell was present in multiple epochs within a day. When plotted, mean firing rate and waveform features generated distinguishable clusters (Extended Data Fig. 5), the boundaries of which were defined as follows: fast-spiking interneurons (FSIs): firing rate >3 Hz, peak width <0.2 ms, and a ratio of trough width to peak width (TPR) <2.7 (TPR was estimated by k-means clustering and was more reliable than exact trough width for FSIs); tonically-active neurons (TANs): <5% of ISIs less than 10 ms, a median ISI >100 ms, and peak width and trough width above the 95^th^ percentile for the remainder of the units; unclassified units had low TPR and/or narrow trough widths (<0.2 or 0.3 ms) but firing rates <2 Hz; all other units were considered putative medium spiny neurons (MSNs). Only MSNs and FSIs are included in the current study.

Hippocampal units were also classified according to mean firing rate and peak and trough width. Putative interneurons were defined as having firing rates >5 Hz, peak width <0.2 ms and trough width <0.3 ms. All other non-unclassified units were considered putative pyramidal cells (used only in Extended Data Fig. 6).

### SWR-triggered spiking activity

For all analyses of SWR-aligned spiking, we created SWR-onset-triggered rasters (1 ms bins) in a 1 s peri-SWR window. From this raster, the mean firing rate was smoothed with a 10 ms s.d., 80 ms wide Gaussian kernel to generate a peri-event time histogram (PETH). For analyses based on z-scored firing rates (e.g. Fig. 2c,g), the raster was padded with a 100x repetition of its start and end values, smoothed, unpadded, and z-scored to the pre-SWR period −500 to 0 ms.

For multiunit activity (MUA) analysis in dH and vH, we thresholded all spike events at 40 µV on tetrodes with clear multiunit firing in the pyramidal layer. In Rats 4 and 5, MUA was extracted by post-hoc thresholding of the 600-6000 Hz filtered LFP. SWR-triggered MUA spike counts were summed across tetrodes and then divided by the total time per bin to calculate a mean firing rate per animal.

To detect significant SWR-modulation of NAc cells, we followed a procedure described previously^40^. Briefly, for each cell, we circularly shuffled each SWR-triggered spike train by a random amount up to ±0.5 s to generate 5000 shuffled PETHs. We then calculated the summed squared difference of the real PETH relative to the mean of the shuffles in a 0-200 ms window post SWR-onset, and compared it to the same value for each shuffle relative to the mean of the shuffles. Significance at p<0.05 indicates that the real modulation exceeded 95% of the shuffles. The direction of modulation was defined from a modulation index, calculated as the mean firing rate in the 0-200 ms window minus the mean baseline firing rate from −500 to −100 ms, divided by the mean baseline firing rate. This sign of this index was used to assign cells as significantly positively or negatively SWR-modulated.

To categorize cells according to both dSWRs and vSWRs, we only included cells that fired at least 50 spikes in the peri-event rasters for both types of SWRs. Cells were subsequently categorized according to their significance and direction as unmodulated (Neither, N), dSWR-significant only (D only), vSWR-significant only (V only), significant during both (Both), dSWR-activated (D+), dSWR-suppressed (D-), vSWR-activated (V+), vSWR-suppressed (V-), or combinations of these: D+V+, D+V-, D-V+, or D-V-. In both wake and sleep, we observed more dSWR- and vSWR-modulated cells than the chance level of 5%. To assess the significance of the “both” modulation categories, we compared each fraction to the chance overlap of our empirical fractions of dSWR- and vSWR-modulated cells using a nonparametric z-test for proportions. We defined “modulation amplitude” as the mean z-scored firing rate of each cell (relative to the pre-SWR period −500 to 0 ms) in the 0-200 ms window following SWR onset.

### Potential duplicate cell control

To control for the possibility that cells stably recorded on the same tetrode across days could have been counted more than once and could influence any of our results, we excluded potential duplicate cells based on waveform similarity^41^ (illustrated in Extended Data Fig. 5). We first established a waveform correlation threshold based on cells recorded on different tetrodes on the same day, which are different cells by definition. For each pair of cells, we aligned their mean waveforms at the peak (on the maximum channel) of one of the cells and calculated a Pearson’s correlation coefficient on each channel (channel 1 of cell A was compared to channel 1 of cell B, and so on). In cases where the waveform snippets were different lengths (due to different spike extraction in Matclust as compared to Mountainsort), we aligned the snippets at their peaks and padded the edges with zeroes as needed. The resulting r values for each channel were then averaged to establish a mean r for that pair. The 95^th^ percentile of r values in this different-cell distribution, 0.979 for wake and 0.980 for sleep, was taken as the threshold for waveform correlation. Next, if a tetrode was moved ≥78 µm across days, we considered the newly acquired cells to be “unique.” If a tetrode was moved less than 78 µm between days, we computed the mean r for all pairs of cells on that tetrode across all previous days of similar depth. This could exclude cells that disappeared and “came back” across multiple days, even though this scenario would seem to be unlikely. Cell pairs with a mean r greater than the threshold were tagged as potential duplicates. We first kept cells from the day with the most cells on that tetrode (randomly selected if multiple days tied for maximum cell count). If a given potential duplicate cell had not yet been kept, one instance of that cell was randomly selected to keep. We note that this system will result in some false positive exclusions and false negative inclusions; different MSNs can have highly similar waveforms even though they are different cells (false positive), and waveforms can change dramatically from day to day even for the same cell due to changes in cell health or relative position of the tetrode (false negative). However, applying this conservative control did not change any of our main results.

### NAc neuron task firing

We analyzed trajectory firing patterns using two methods: normalized trial time and linearized position. In the normalized trial time method, each trial was split into normalized progression of time spent at the well (from nosepoke to when the animal turns around; “well”) and time spent moving along the path between wells (from turnaround to next nosepoke; “path”). The turnaround time was detected by a >4 cm movement in the x direction, a change in head direction of >0.25 radians (∼14°), and a speed of >2 cm/s. Additionally, we required that the animal had moved away from the well in the y-direction one second in the future, otherwise the turnaround time was incremented. Path and well time were divided into bins of 0.5% of the total completion time. Firing rate was calculated by dividing the number of spikes in each bin by the bin width in seconds on that particular trial (excluding spikes during either dSWRs or vSWRs), smoothing the rate with a 5 bin (2.5%) s.d., 40 bin (20%) wide Gaussian kernel, and then averaging across trials of the same trajectory type (defined by start and end well). We further attempted to control for variation in the animal’s behavior on individual trials in three ways: by only calculating mean trajectory rates when there were at least 3 trials on that trajectory; by performing a pairwise speed profile correlation across trials and only accepting trials that fell at or above the 25^th^ percentile of speed similarity values; and by only accepting trials with a duration at or below the 75^th^ percentile of the trial length distribution. These methods excluded trials that were long, slow, or had many stops.

In the linearized position method, we projected the animal’s 2D position to a line connecting each junction and endpoint of the maze, generating a linearized position relative to the start of each trajectory defined by start and end well. Each trajectory thus contained a specific set of maze segments, and we again controlled for behavioral variation by only accepting trials where the animal deviated ≤12 linear cm onto segments not included in the current trajectory (this allowed for small “head swings” onto neighboring segments). From the set of included trials on each trajectory, we calculated a firing rate per linear position bin of 2 cm (divided by total time occupancy in that bin), smoothed it with a 2 cm s.d., 10 cm wide Gaussian kernel, and calculated the mean rate on that trajectory within day. Trajectories missing more than 5 linear position samples (as a result of diode occlusion or due to low time occupancy) were excluded from firing similarity analysis (5 or fewer missing samples were interpolated), and linearized distance was normalized before pairwise correlation across trajectories (below).

To assess the firing similarity of a given cell across trajectories that differ in spatial location and direction, we focused on the 6 rewarded trajectories (across SA and SB) depicted in Fig. 3a. We calculated the coefficient of determination between the mean firing profiles of each pair of trajectories (as a function of normalized trial time or linearized position), and then took the mean r^2^ across pairs. We controlled for the effect of firing rate by matching cells in the V+ population to D+ and N cells with the closest firing rates, generating subsampled D+ and N populations. Note that the variety of behavioral controls applied to both methods excluded slightly different numbers of cells, depending on whether the cells were active on enough trials that passed our criteria to compute an r^2^.

We explored a variety of additional task-related firing parameters, reported in Extended Data Fig. 9. Trial-by-trial correlation was performed with the same controls for behavioral variability as described above. Specifically, we correlated pairs of individual trials on the same trajectory to get a mean r^2^ for that trajectory, and then took the mean r^2^ across trajectories. In the linearized position version, this was done with spike counts in 5 cm bins as opposed to smoothed firing rate, as time occupancy in any given bin could be very low on a given trial. Spatial information^42^ was calculated by treating each maze segment (Extended Data Fig. 9c) or each vertical maze segment plus direction (Extended Data Fig. 9d) as a single spatial bin. Path vs. well preference was calculated from each cell’s mean path and well (excluding SWRs) firing rates across trajectories in normalized trial time, as (path – well)/(path + well), such that values greater than zero indicate path preference and values less than zero indicate well preference. Trajectory direction preference was calculated from linearized position, as the absolute area between leftward and rightward trajectory firing rate curves on the same maze segments, divided by their sum (values closer to 1 indicate a stronger preference in one direction, either left or right). Toward vs. away-from-well preference measures the preference of a cell for path movement toward or away from wells (regardless of reward outcome), and is calculated from the linearized position firing rate curves on the same vertical segment in opposite directions (i.e. the area between the two curves divided by their sum).

### NAc neuron reward and reward history firing

To calculate reward vs. error preference at the wells (based on current reward or error), we used two methods. In the first method (Extended Data Fig. 9h,i), we calculated firing rate on rewarded and error well visits as a function of normalized time at the wells (0.5% bins, excluding SWR spikes, smoothed with a 1.5% s.d., 12% wide Gaussian kernel), again applying a pairwise speed profile correlation to only include trials that fell at or above the 25^th^ percentile of speed similarity values. We only included cells for which at least 3 trials of both types met the above criteria. We then calculated a reward vs. error index (RI) per cell which equaled the area between the mean rewarded firing rate and mean error firing rate curves divided by their sum, exclusively in time bins for which the mean speed on both rewarded and error trials was <4 cm/s. Significance of reward preference (>0) or error preference (<0) was calculated by a one-tailed permutation test from the set of rewarded and error trials.

In the second method for reward preference at the wells (Extended Data Fig. 9j,k), we calculated rewarded and error firing rates as a function of time from nosepoke (0 to 4 s, 100 ms bins). We then computed a reward vs. error index (RI) as the difference in mean rewarded and error firing rates post-reward-delivery time (2 to 4 s, excluding SWR spikes and times), divided by their sum. We again controlled for speed by excluding any trial where the animal spent more than 5 position samples (150 ms) moving faster than 4 cm/s, and required at least 5 included trials of both types to compute an RI. Significance was again assessed with a one-tailed permutation test in the 2-4 s window.

To examine reward history preference, we calculated mean firing rate on normalized path time using the same methods as for trajectory firing (but smoothed with a 1.5% s.d., 12% wide Gaussian kernel), now comparing paths following a rewarded well visit or an error well visit (trials are defined by reward outcome of trial n-1). We only included cells for which at least 3 trials of both types met our speed and trial length criteria. The reward history preference was calculated as area between the curves divided by their sum and tested for significance of reward preference (>0) vs. error preference (<0) with a one-tailed permutation test.

### Spike cross-correlations

Spike cross-correlations between pairs of D+ or V+ NAc MSNs were calculated in 10 ms bins at up to 0.5 s lag. Each CCH was first normalized by the square root of the product of the number of spikes from each cell. To z-score the CCH of each cell pair, one of the spike trains was circularly shuffled 1000 times (by a random amount up to ±half the mean immobility period length) to create 1000 shuffled CCHs. Each real and shuffled CCH was smoothed with a 20 ms s.d., 160 ms wide Gaussian kernel. The real cross-correlation values were then z-scored relative to the distribution of shuffled values within each bin. We averaged the smoothed cross-correlation z-score in a 20 ms bin around 0 (±10 ms) to get an approximate “0 lag” value, and the distribution of z-scores at 0 lag across pairs was converted to a cumulative density function (Fig. 4).

### Data reporting

No statistical methods were used to predetermine sample size. The minimum number of required animals was established beforehand as four or more, in line with similar studies in which this number yields data with sufficient statistical power.

### Data availability

Data are available from the authors upon reasonable request.

### Statistics

All statistical tests were two-sided unless otherwise specified.

### Code availability

All custom code is available from the authors upon reasonable request.

---

**Extended Data Figure 1.**
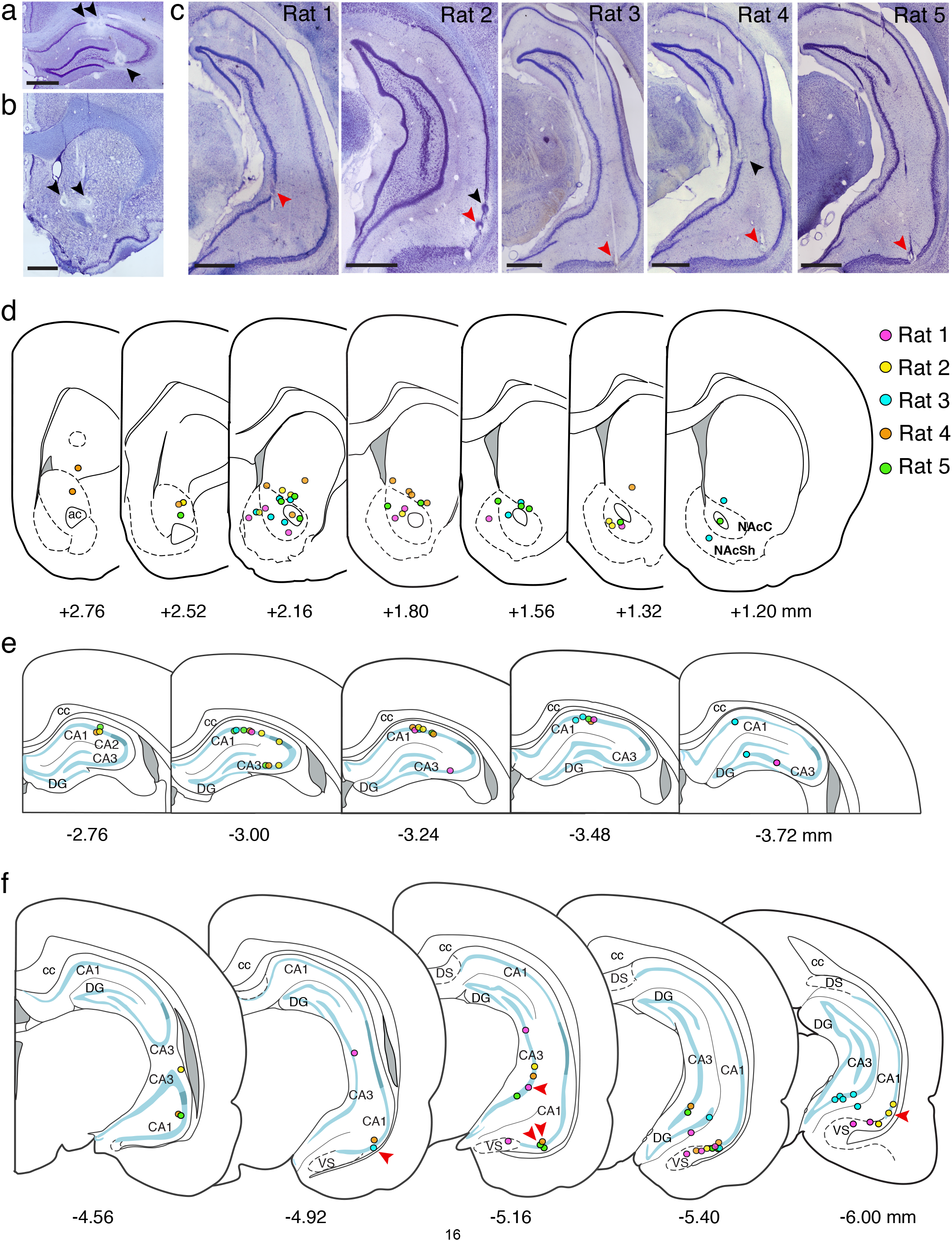
Histological identification of recording sites. **a, b**, Nissl-stained coronal sections showing example tetrode lesions marked by black arrows in (**a**) dCA1 and dCA3 and (**b**) NAc, in Rat 5. **c**, Nissl-stained coronal sections showing example tetrode lesions in vH. Each section includes the tetrode used for vSWR detection in each rat (red arrows). In **a**, **b**, and **c**, scale bars are 1 mm. **d**, Summary of all NAc recording locations across rats, aligned to the nearest representative section adapted from the Paxinos & Watson Wistar rat brain atlas (2007). Dots mark the last recorded depth of each tetrode at the end of the experiment. Dotted lines indicate approximate borders of NAc core and shell. Grey shaded regions represent ventricles. All distances are in AP coordinates (mm) relative to Bregma and correspond to original plate labels from the atlas; note that theses distances do not correspond exactly to true recording locations in our Long-Evans rats and are for illustration purposes only. For true implant coordinates, see Online Methods. **e, f**, Summary of all recording sites in dH (**e**) and vH (**f**). Light blue regions represent stratum pyramidale of CA1/CA3 or granule cell layer of dentate gyrus, darker blue regions represent stratum pyramidale of CA2. Grey shaded regions represent ventricles. Red arrows indicate the 5 sites used for vSWR detection (1 site per animal). Abbreviations: ac, anterior commissure; NAcC, nucleus accumbens core; NAcSh, nucleus accumbens shell; cc, corpus callosum; DG, dentate gyrus; DS, dorsal subiculum; VS, ventral subiculum.

**Extended Data Figure 2.**
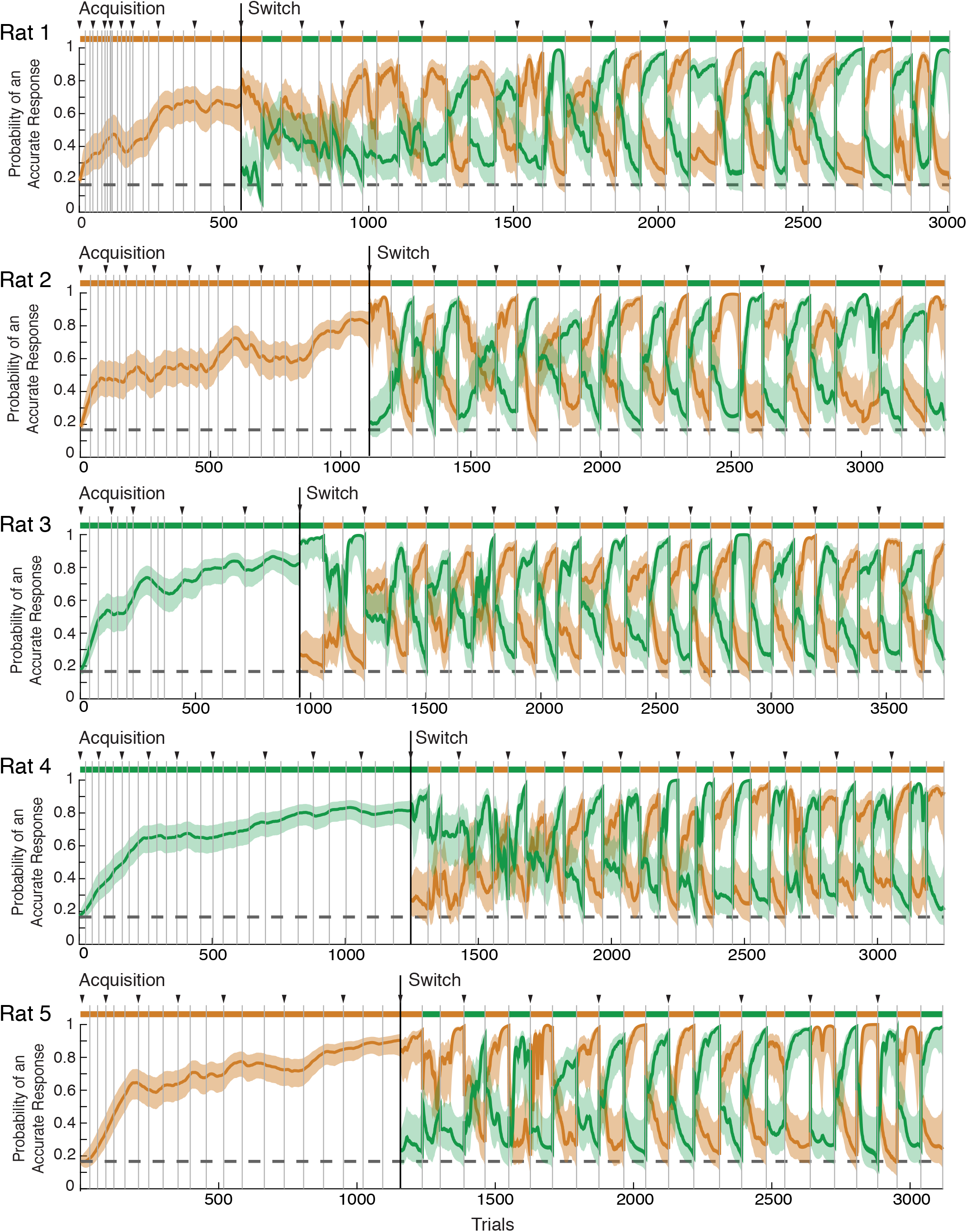
Multiple-W behavior. Behavior from each rat expressed as the probability that the rat is making an accurate choice on each trial according to Sequence A (orange) or B (green) (see Online Methods). Solid line indicates the mode of the probability distribution, shaded region indicates the 90% confidence interval. Colored bars at the top of each plot indicate the rewarded sequence, grey vertical lines mark epoch boundaries, black triangles mark the start of days, and the horizontal dotted line indicates chance performance: 1/6 (0.167).

**Extended Data Figure 3.**
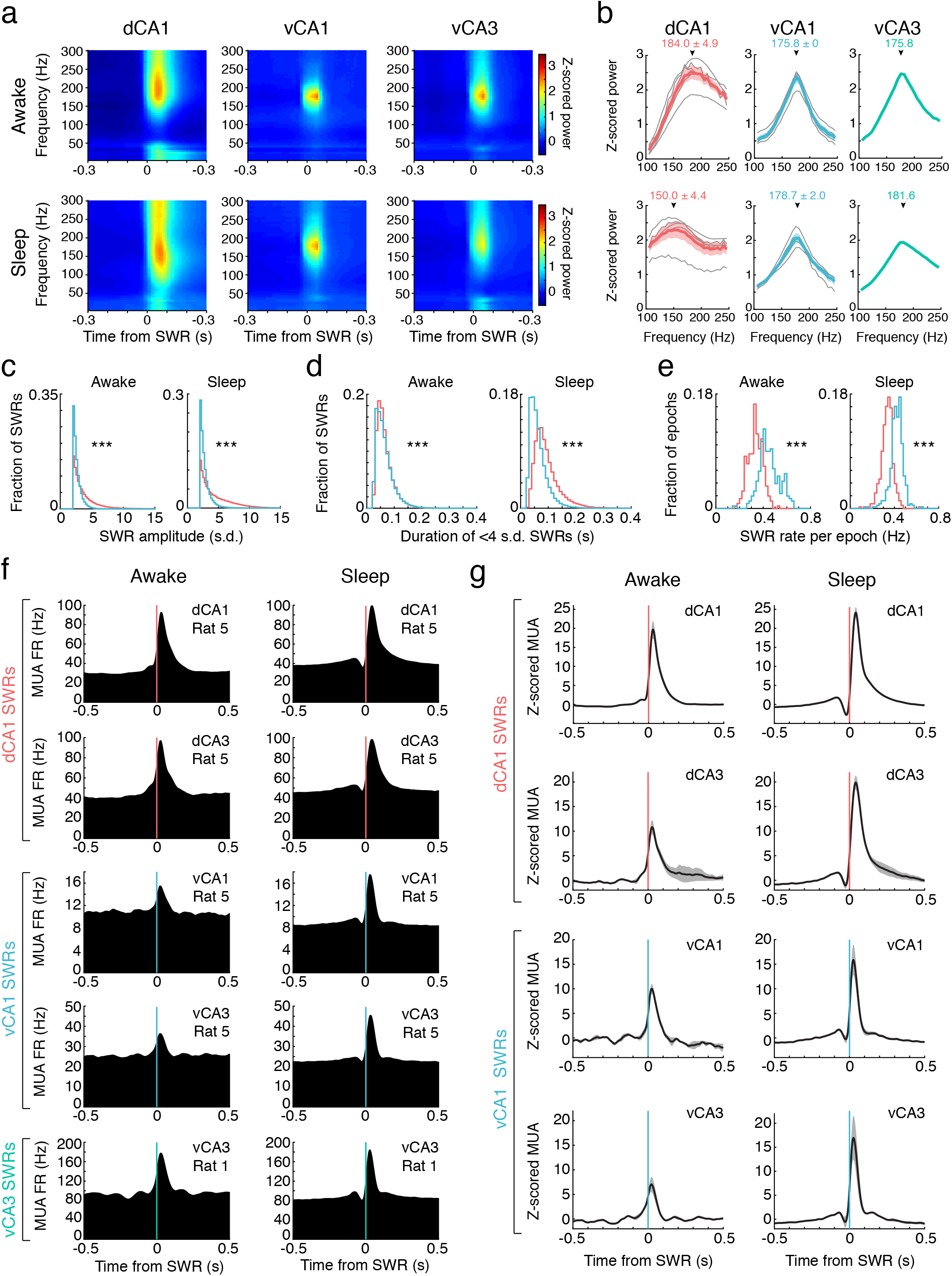
Characterization of dH and vH SWRs. **a,** Examples of mean SWR-onset-triggered spectrograms on one tetrode in one rat, from each detection region, during awake immobility (top) and sleep (bottom). Left: all dCA1 SWRs (n=9,642 awake, 26,048 sleep) in Rat 5; Center: all vCA1 SWRs (n=9,672 awake, 31,553 sleep) in Rat 5; Right: all vCA3 SWRs (n=18,798 awake, 24,536 sleep) in Rat 1 (the only rat in which vCA3 SWRs were used). All SWRs >2 s.d. are included. Color bar shows power, z-scored within each epoch and averaged across epochs and days. In **a** and **b**, SWRs were detected as described in Online Methods (3 dCA1 tetrodes and 1 vH tetrode per animal). **b**, Mean z-scored power (a slice of the SWR-triggered spectrogram) at frequencies between 100-250 Hz, at the time of peak ripple power for each animal during awake immobility (top row) and sleep (bottom row). Grey curves indicate individual animals, colored curves indicate mean ± s.e.m. across animals. Arrows indicate mean ± s.e.m. peak ripple frequency. Left: dCA1 SWRs (n=51,333 awake, 115,158 sleep, 5 rats); Center: vCA1 SWRs (n=47,222 awake, 120,271 sleep, 4 rats); Right: vCA3 SWRs (n=18,798 awake, 24,536 sleep, 1 rat). As vCA1 and vCA3 SWRs occurred at nearly the same frequency, we pooled these ventral SWRs (vSWRs) for the remainder of the study. **c**, Distributions of SWR amplitudes across animals. In both wake and sleep, dSWRs (pink) are typically larger amplitude than vSWRs (blue; ***p=0, Wilcoxon rank-sum test), consistent with previous work^20^. In **c**, **d**, and **e**, only one dCA1 tetrode per animal was used for SWR detection to allow for direct comparison to vSWRs. (in **c** and **e**, awake n=54,496 dSWRs, 74,355 vSWRs; sleep n=118,897 dSWRs, 146,995 vSWRs). **d**, Distributions of SWR durations for <4 s.d. SWRs, across animals. In both wake and sleep, dSWRs (pink) are significantly longer than vSWRs (blue), although this difference is more pronounced in sleep (***p<10^40^, Wilcoxon rank-sum test; awake n=38,170 dSWRs, 71,029 vSWRs; sleep n=66,382 dSWRs, 135,035 vSWRs). **e**, Distributions of dSWR (pink) and vSWR (blue) rate per epoch, expressed as the fraction of epochs with each rate (out of 256 awake epochs and 411 sleep epochs across animals). For awake epochs on the maze, rate was calculated per total time spent at <4 cm/s. For sleep epochs, rate was calculated per time spent in NREM sleep (see Online Methods). In both wake and sleep, the vSWR rate is typically higher than the dSWR rate (***p<10^40^, Wilcoxon rank-sum test). **f**, Examples of hippocampal multiunit activity (MUA) in wake and sleep, aligned to the onset of dSWRs (pink lines), vCA1 SWRs (blue lines), or vCA3 SWRs (aqua lines). In the upper right of each panel is the region and rat from which MUA was detected. Firing rate was calculated from the summed spike count from all tetrodes in the region. **g**, Mean z-scored multiunit firing rate across animals. Grey shaded region indicates s.e.m. Top two rows: dCA1 and dCA3 MUA aligned to dCA1 SWRs (n=5 rats). Bottom two rows: vCA1 MUA (n=4 rats) and vCA3 MUA (n=3 rats) aligned to vCA1 SWRs. Z-scores were calculated within animal relative to the pre-SWR period (−500 to 0 ms). Strong similarity in activation timing of vCA3 MUA between vCA1 SWRs and vCA3 SWRs also provided support for the use of vCA3 for SWR detection in Rat 1.

**Extended Data Figure 4.**
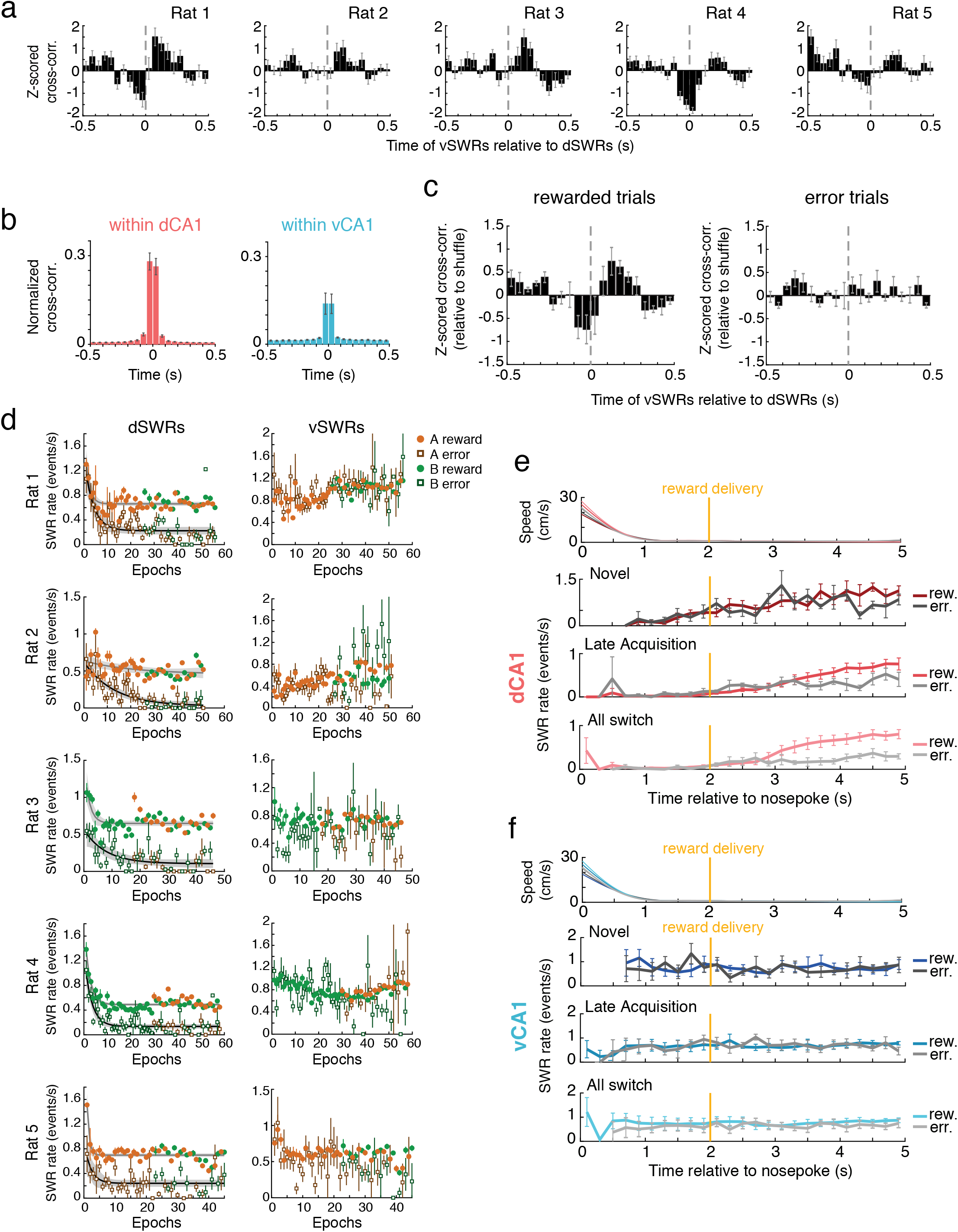
Awake SWR asynchrony and timing relative to reward. **a**, Cross-correlation histograms (CCH) of awake vSWRs relative to dSWRs within each animal, z-scored relative to shuffle. Note a lack of synchrony at less than 50 ms lag across animals. Error bars indicate s.e.m. across days (n=15-19 days) for each animal. **b**, Mean CCH showing high synchrony of awake SWRs between tetrodes within dCA1 (left, n=5 rats) or vCA1 (right, n=3 rats with >1 tetrode in vCA1), normalized to the number of SWRs on one tetrode. Complementary to Fig. 1f. Error bars indicate s.e.m across animals. **c**, Asynchrony between dSWRs and vSWRs is present on both rewarded (left) and error (right) trials. A minimum of 3 s immobility following the time of reward delivery (or expected reward delivery on error trials) was required to include a trial’s SWRs (n=5 rats, error bars are s.e.m.). **d**, Changes in rewarded vs. error SWR rate over task exposure, as a function of maze epochs per animal. Filled circles indicate mean SWR rate across rewarded trials (well visits) within the epoch (± s.e.m.), open squares indicate mean SWR rate across error trials. Each point is color coded by the rewarded sequence of that epoch (Sequence A: orange; Sequence B: green). Left: dSWRs; black line indicates an exponential fit ± 95% confidence interval on error SWR rate, grey line indicates exponential fit on rewarded SWR rate. Right: vSWRs; note that some error bars extend above the y-axis limit. **e**, Timing of dSWRs relative to reward delivery, by behavioral stage. Gold line in **e** and **f** indicates reward delivery or its expected time on error trials, 2 s after nosepoke at the well. Top: mean speed in each behavioral stage; note that each rat’s head takes ∼1 s to fully decelerate and dip into the reward well. Below: mean SWR rate across animals in 200 ms bins for rewarded and error trials during the first 100 trials of Multiple-W behavior (“Novel”), trials occurring at >0.6 probability correct on the first sequence (“Late Acquisition”), and during all Switch epochs (“All switch”). Shades of red are rewarded trials, shades of grey are error trials. **f**, Timing of vSWRs relative to reward delivery, by behavioral stage, similar to **e**. Speed data at top are repeated from **e**. Note that since reward rate is only calculated when at least 2 s of data are present below 4 cm/s in each bin, empty bins indicate too little data to calculate an SWR rate. Shades of blue are rewarded trials, shades of grey are error trials.

**Extended Data Figure 5.**
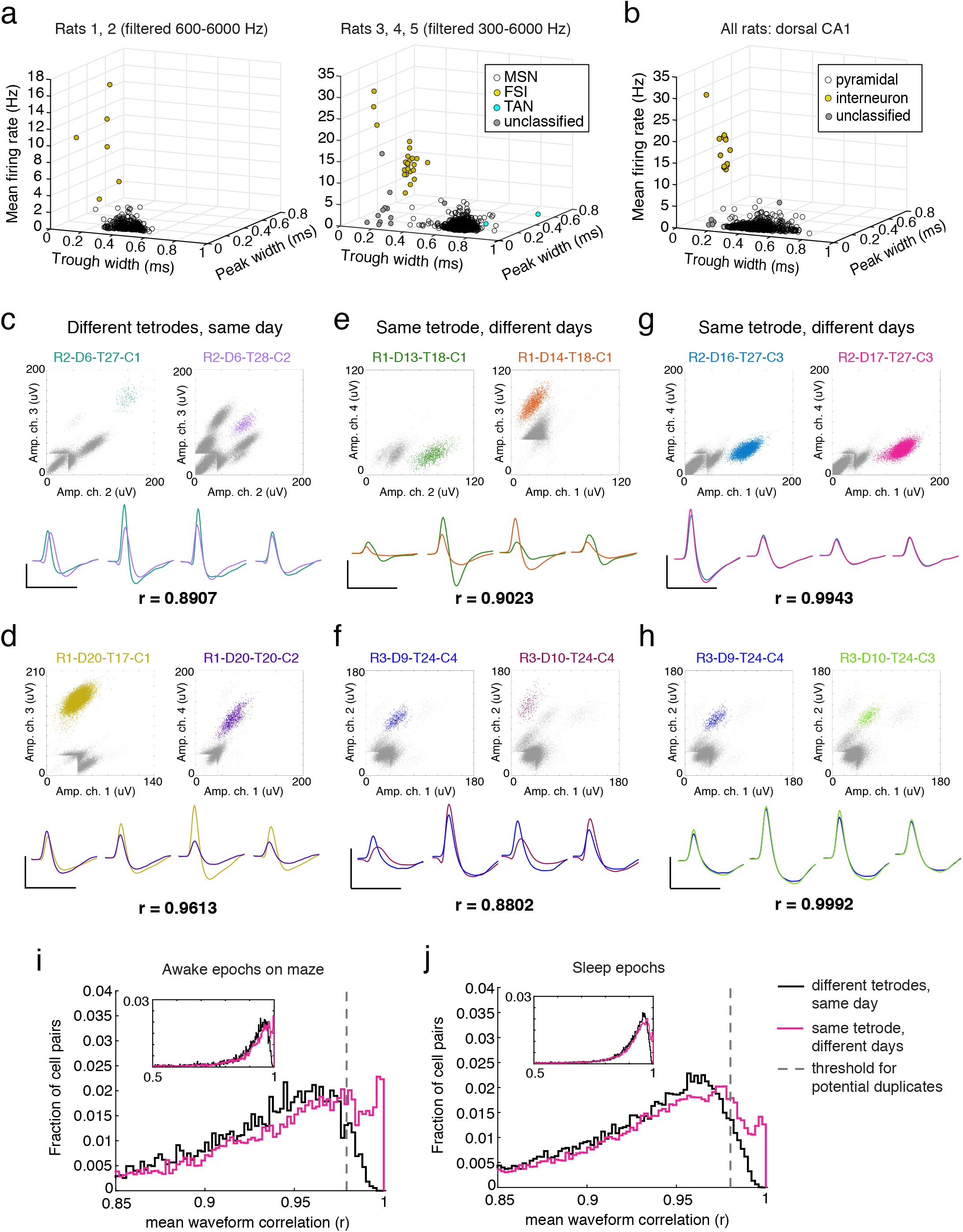
NAc cell type classification and waveform correlation control. **a**, Classification of putative NAc cell types by waveform properties and mean firing rate. Each point represents a single unit. Number of units (wake and sleep included): MSNs: 1799, FSIs: 30, TANs: 2, unclassified: 12. Data from Rats 1 and 2 (left) were classified separately from Rats 3-5 (right) due to narrower bandpass filtering at time of acquisition, as this changes waveform shape. **b**, Classification of putative hippocampal cell types in dCA1 (contributes to Extended Data Fig. 6). Number of units (wake and sleep included): pyramidal: 1218, interneuron: 13, unclassified: 18. **c-h**, Illustration of waveform correlation metric to assess cells as “unique” or “duplicate.” Each panel shows an example pair of cells with their mean Pearson’s r value across all 4 channels listed at the bottom. Top row: each cell’s cluster in amplitude space on two channels of the tetrode; each point is the maximum amplitude of a spike. Colored spikes represent the chosen unit; grey spikes represent all other threshold crossings. Cell ID code lists the rat number, day number (since beginning of data acquisition), tetrode number, and cell number (assigned out of the total cells on that tetrode, for that day). Bottom row: mean waveforms of the cell pair on each channel, aligned to the first cell’s peak on its maximum channel. All scale bars are 60 µV on the y-axis, 1 ms on the x-axis. **c and d**, Cell pairs which were recorded on different tetrodes on the same day, thus by definition are different cells. **e and f**, Cell pairs recorded on the same tetrode on consecutive days at the same depth, but which have an r below the threshold for a duplicate cell, indicating that they are likely different cells. **g and h**, Cell pairs recorded on the same tetrode on consecutive days at the same depth with an r above the threshold, and thus are classified as potential duplicates across days. The left (blue) cell in **h** is the same cell on the left in **f**, and the clustering projection on the right in **h** is the same as the right projection in **f**. **i**, Distributions of mean waveform r for all cell pairs, calculated from awake epochs on the maze. Black: pairs on different tetrodes on the same day. Magenta: pairs on the same tetrode at similar depths across days. Inset shows zoomed-out view of the whole distribution. Vertical dotted line shows the threshold used to classify a cell pair as potential duplicates, r=0.979, which is the 95^th^ percentile of the black distribution. **j**, Similar to **i**, but for waveform r calculated from sleep epochs. Sleep threshold: r=0.980. Cells present in both wake and sleep were classified as potential duplicates based on their r in wake.

**Extended Data Figure 6.**
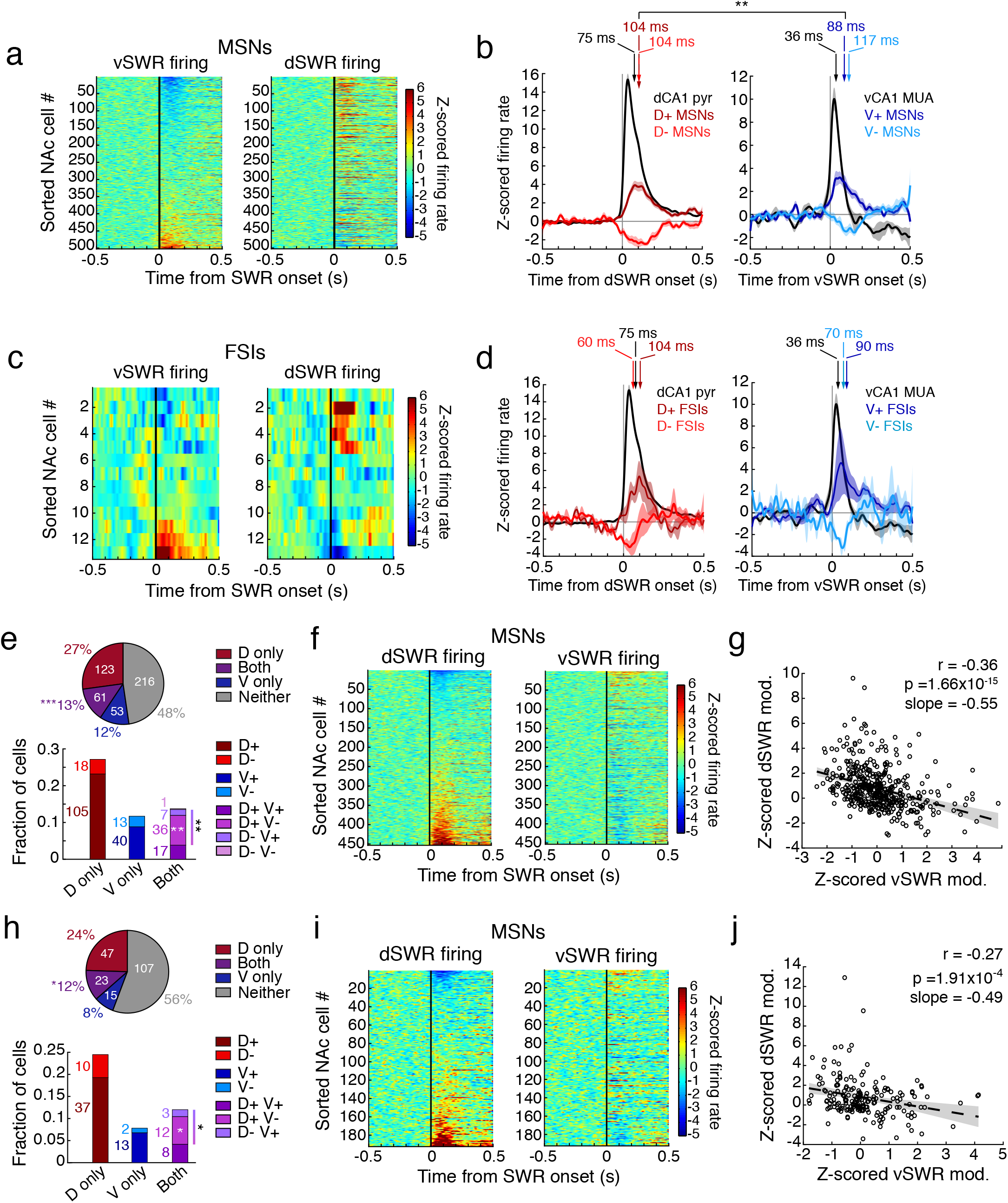
NAc modulation during awake SWRs. **a**, Opposing modulation is not a result of cell ordering. Left: vSWR-aligned z-scored PETHs for each MSN ordered by its modulation amplitude (mean z-scored firing rate in the 200 ms following SWR onset). Right: dSWR-aligned z-scored PETHs for the same ordered MSNs shown on the left. Same MSNs as shown in Fig. 2c. **b**, Timing of significantly modulated MSN ensemble activity during awake dSWRs (left) and vSWRs (right), overlaid on dCA1 pyramidal cell firing (n=332 cells) during dSWRs (left) and vCA1 multiunit activity (n=4 rats) during vSWRs (right). Each curve represents the mean ± s.e.m. z-scored firing rate of the hippocampal (black), D+ (n=49 cells, dark red), V+ (n=16 cells, dark blue), D- (n=13 cells, light red), or V- (n=14 cells, light blue) populations. MSN subpopulations here are restricted to cells that passed the potential duplicate cell control. Arrows indicate the center of mass in the 0-200 ms post-SWR interval of each subpopulation’s activity. The D+ center of mass occurs significantly later than the V+ center of mass (**p<0.01, one-tailed permutation test). Timing difference of centers of mass of the D- and V-populations was not significantly different at the p<0.05 level. Also note that the centers of mass of the activated NAc ensembles lag the local activation of dH and vH neurons. **c**, Similar to **a** but for FSIs shown in Fig. 2g. **d**, Timing of significantly modulated FSI ensemble activity during awake dSWRs (left), and vSWRs (right), overlaid on the same hippocampal activity as shown in **b**. Arrows indicate the center of mass in the 0-200 ms post-SWR interval of each population’s activity; no significant differences in timing (permutation tests). **e-g**, Opposing MSN modulation cannot be accounted for by occasional co-occurrence of dSWRs and vSWRs (control for Fig. 2b-d). We removed dSWRs and vSWRs occurring within 250 ms of each other to examine SWRs that occurred in isolation. Fraction modulated during Both: ***p=9.13×10^4^, fraction opposing: **p=0.0046, fraction D+V-: **p=0.0089 (z-tests for proportions). **h-j**, Opposing MSN modulation cannot be accounted for by inclusion of potential duplicate cells (control for Fig. 2b-d). We excluded potential duplicates using the method illustrated in Extended Data Fig. 5. Fraction modulated during Both: *p=0.016, fraction opposing: *p=0.038, fraction D+V-: *p=0.046 (z-tests for proportions).

**Extended Data Figure 7.**
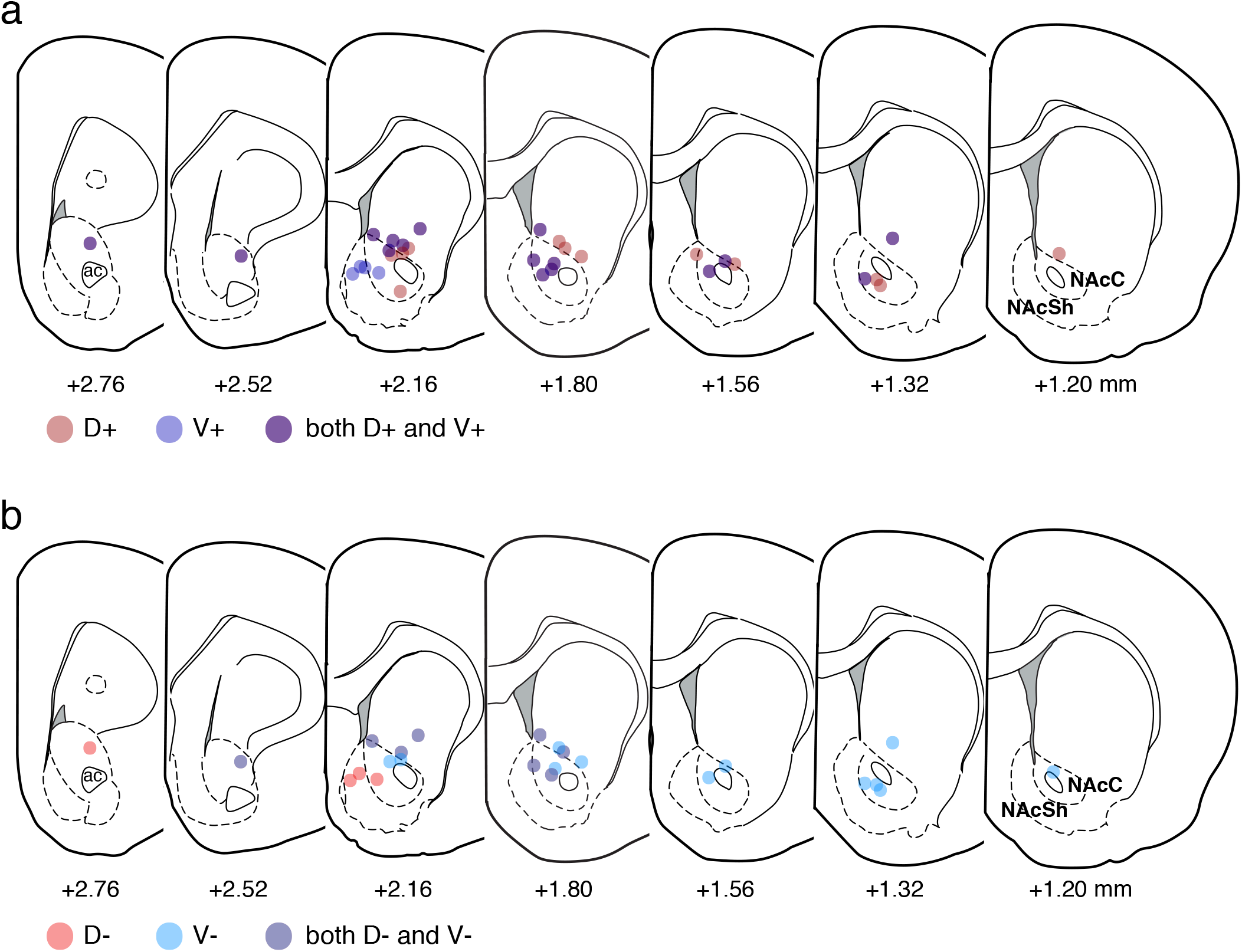
Anatomical locations of NAc neurons modulated during awake SWRs. **a**, Mapping of approximate tetrode locations where at least one D+ (dark red) or one V+ (dark blue) neuron (either MSN or FSI) was detected. All AP coordinates are in mm relative to Bregma and correspond to plate labels from the atlas (see Extended Data Fig. 1). Overlapping colors indicate locations where both D+ and V+ were detected, typically not in the same neuron. Note that borders of core (NAcC) and shell (NAcSh) are approximate, but that D+ modulation is limited largely to the dorsolateral core and its dorsal boundary, whereas V+ modulation is seen in the medial shell and also extends into the core. **b**, Mapping of approximate tetrode locations where at least one D-neuron (light red) or one V-neuron (light blue) was detected, again typically not in the same neuron. Note coincidence of D-sites with V+ sites and V-sites with D+ sites.

**Extended Data Figure 8.**
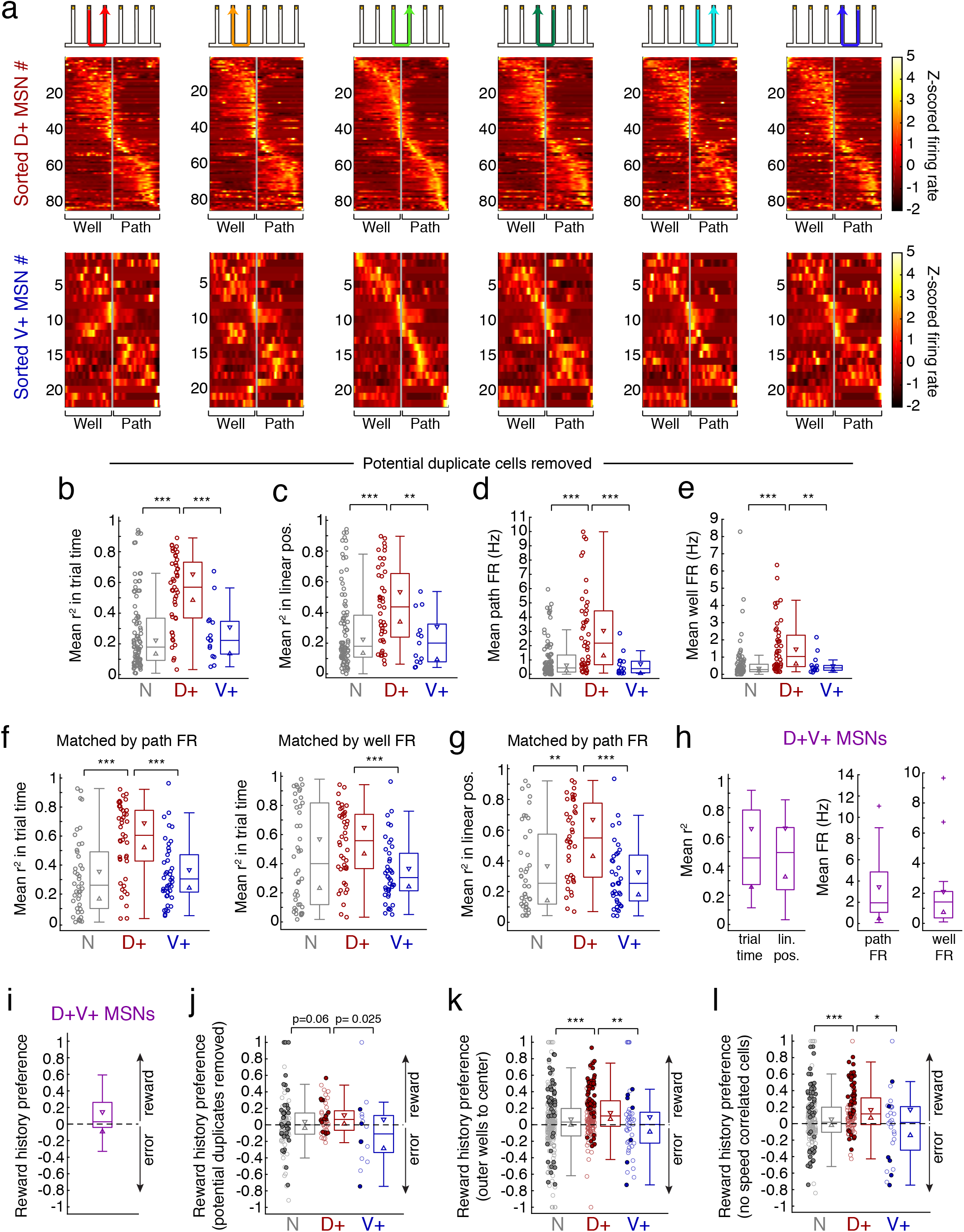
Population task-related firing patterns of D+ and V+ MSNs. **a**, Heterogeneity of trajectory stages represented by NAc MSNs, as a function of normalized trial time. Specific trajectories are shown in cartoons above the heat plots. D+ MSNs (top) and V+ MSNs (bottom) are sorted by their peak z-scored firing location on the third trajectory (light green). Z-scores are calculated within cell from the mean firing rate profile on each trajectory. Only cells which were active on enough trials (see Online Methods) of all six trajectories are shown here, such that these cells predominantly correspond to the Switch phase of the task (D+ n=85 cells, V+ n=22 cells). **b-e**, Controls for Fig. 3c-f with potential duplicate cells removed using the method illustrated in Extended Data Fig. 5. In **b, d, e**: N n=93 cells, D+ n=45 cells, V+ n=15 cells; in **c**: N n=90 cells, D+ n=44 cells, V+ n=13 cells. **p<0.01, ***p<0.001. All tests between populations in **b-l** are Wilcoxon rank-sum tests with Bonferroni correction for multiple comparisons, setting significance level at p<0.017. **f**, D+ MSN firing pattern similarity across trajectories in normalized trial time is maintained when populations are matched for mean firing rate on the path (left: n=42 cells in each subpopulation; N vs. D+: ***p=1.18×10^4^; D+ vs. V+: ***p=1.22×10^4^) or well (right: N vs. D+: p=0.34; D+ vs. V+: ***p=4.17×10^4^). **g**, Similar to **f** but for firing similarity in linearized position, matched by mean firing rate on the path (n=40 cells in each subpopulation; N vs. D+: **p=0.0016; D+ vs. V+: ***p=2.55×10^5^). **h**, D+V+ MSNs resemble D+ MSNs (see Fig. 3c-f) in firing pattern similarity across trajectories and in firing rate. All D+V+ MSNs are included. Crosses indicate outliers. **i**, Reward history preference of all D+V+ MSNs is not significantly greater than zero (p=0.11, one-tailed signed-rank test). **j**, Control for Fig. 3h with potential duplicate cells removed. N n=104 cells, D+ n=49 cells, V+ n=16 cells. D+ population is still significantly shifted greater than zero (p=0.0041, one-tailed signed-rank test). **k**, Control for Fig. 3h, examining only paths coming from the outer two wells to the center well of each rewarded sequence. This aims to control both for upcoming choice and reward expectation, as returns to the center well of each sequence are always rewarded. D+ population is significantly shifted greater than zero (p=3.51×10^11^, one-tailed signed-rank test). N (n=228 cells) vs. D+ (n=159 cells): ***p=4.87×10^5^; D+ vs V+ (n=43 cells): **p=0.0042. **l**, Control for Fig. 3h removing cells that are significantly correlated with trial-by-trial changes in in mean speed. This aims to control for the possibility that differing speed on trials following reward vs. error could account for reward history preference. D+ population is significantly shifted greater than zero (p=9.67×10^9^, one-tailed signed-rank test). N (n=172 cells) vs. D+ (n=107 cells): ***p=6.68×10^4^; D+ vs. V+ (n=28 cells): *p=0.013.

**Extended Data Figure 9.**
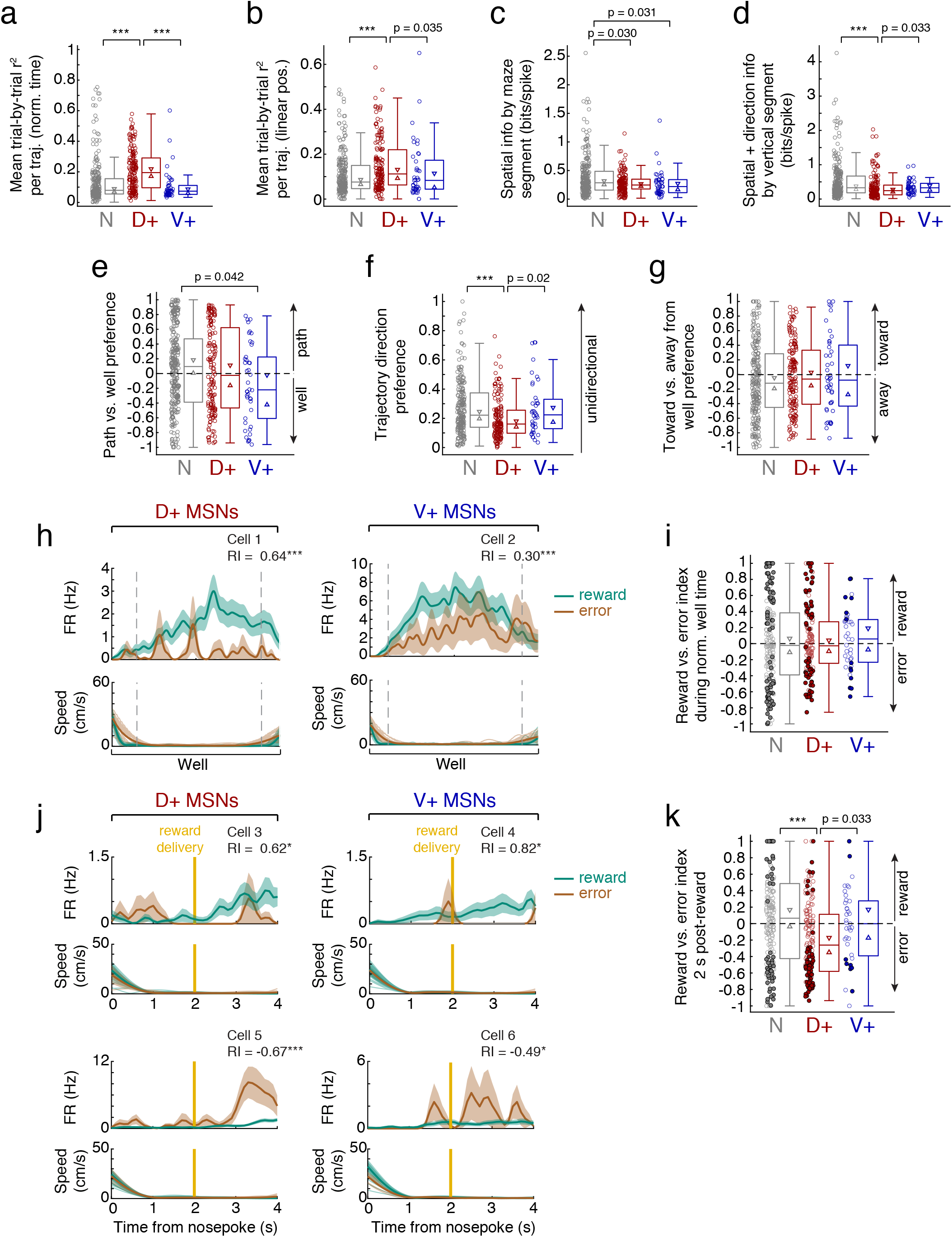
Task-related firing pattern parameters explored for D+ and V+ MSNs. **a**, Mean trial-by-trial correlation, calculated across individual traversals of each trajectory in normalized trial time and then averaged across trajectories to get a single r^2^. Note that the V+ and N populations have lower trial-by-trial similarity (are more variable) than the D+ population. N (n=235 cells) vs. D+ (n=159 cells): ***p=2.14×10^12^; D+ vs. V+ (n=45 cells): ***p=1.09×10^7^. All tests between populations in **a-g, i, k** are Wilcoxon rank-sum tests with Bonferroni correction for multiple comparisons, setting significance level at p<0.017. **b**, Mean trial-by-trial correlation in linearized position, based on spike counts in 5 cm bins. Again, note lower similarity of the N (n=225 cells) and V+ (n=41 cells) populations compared to the D+ population (n=159 cells; N vs. D+: ***p=3.63×10^4^). **c**, Spatial information of MSN spiking, where each of 11 maze segments (6 vertical, 5 horizontal) is considered a spatial bin. Both the D+ (n=161 cells) and V+ (n=45 cells) populations trend towards being less spatially-specific than the N (n=246 cells) population. **d**, Spatial and directional information of MSN spiking, similar to **c**, here examining the 6 vertical segments in both directions for 12 total bins. Note significant or trending lower spatial and directional selectivity of the D+ (n=161 cells) compared to the N (n=246 cells; N vs. D+: ***p=3.50×10^5^) and V+ (n=45 cells) populations, consistent with greater generalization across trajectories. **e**, Preference for the path (>0) vs. well (<0) components of all trajectories. While individual cells exhibit strong preferences, the populations do not (D+ n=154 cells, N n=226 cells, V+ n=42 cells). **f**, Trajectory direction preference, where 0 indicates bidirectionality and values closer to 1 indicate stronger unidirectionality for either leftward- or rightward-moving trajectories. Note significant or trending lower directionality of the D+ population (n=152 cells) compared to the N (n=221 cells; N vs. D+: ***p=3.16×10^5^) and V+ (n=40 cells) populations. **g**, Preference for moving toward or away from a reward well (regardless of reward outcome). Values greater than 0 indicate preference for approaching a well, values less than 0 indicate preference for leaving a well. While individual cells exhibit strong preferences, the populations do not (D+ n=161 cells, N n=241 cells, V+ n=44 cells). **h**, Examples of individual MSNs (D+, left; V+, right) showing strong preference for rewarded well visits (teal) compared to error visits (brown) as a function of normalized well time. Top: firing rate (mean ± s.e.m. across trials). Reward vs. error index (RI) is shown in the upper right of each panel (***p<0.001, one-tailed permutation test). Bottom: faded lines indicate speed profiles of individual well visits, thick lines indicate mean speeds. Dotted grey lines flank the time period analyzed for significance, when both rewarded and error mean speeds are <4 cm/s. **i**, Distributions of reward vs. error index during normalized well time, by SWR-modulation category. Values greater than 0 signify higher firing rates when rewarded, values less than 0 signify higher firing rates on errors. Filled circles indicate significantly reward- or error-preferring cells, open circles indicate non-significant cells (D+ n=141 cells, N n=196 cells, V+ n=39 cells). **j**, Examples of individual MSNs (D+, left; V+, right) firing as a function of time since nosepoke. Top row: higher firing rates (mean ± s.e.m. across trials) on rewarded trials compared to error trials in the 2 s following reward delivery time, with individual and mean speeds for these trials shown below. Bottom row: higher firing rates on error trials compared to rewarded trials. RI is shown in the upper right of each cell (*p<0.05, ***p<0.001, one-tailed permutation test in the 2-4 s window). Gold vertical line marks reward delivery (or time of expected reward delivery on error trials). **k**, Distributions of reward vs. error index during 2 s following reward delivery time, by SWR-modulation category (D+ n=147 cells, N n=195 cells, V+ n=37 cells). The D+ population is significantly shifted negative of zero by this metric (p=5.15×10^7^, one-tailed signed-rank test) and shows significantly less post-reward preference than the N population (***p=9.06×10^6^).

**Extended Data Figure 10.**
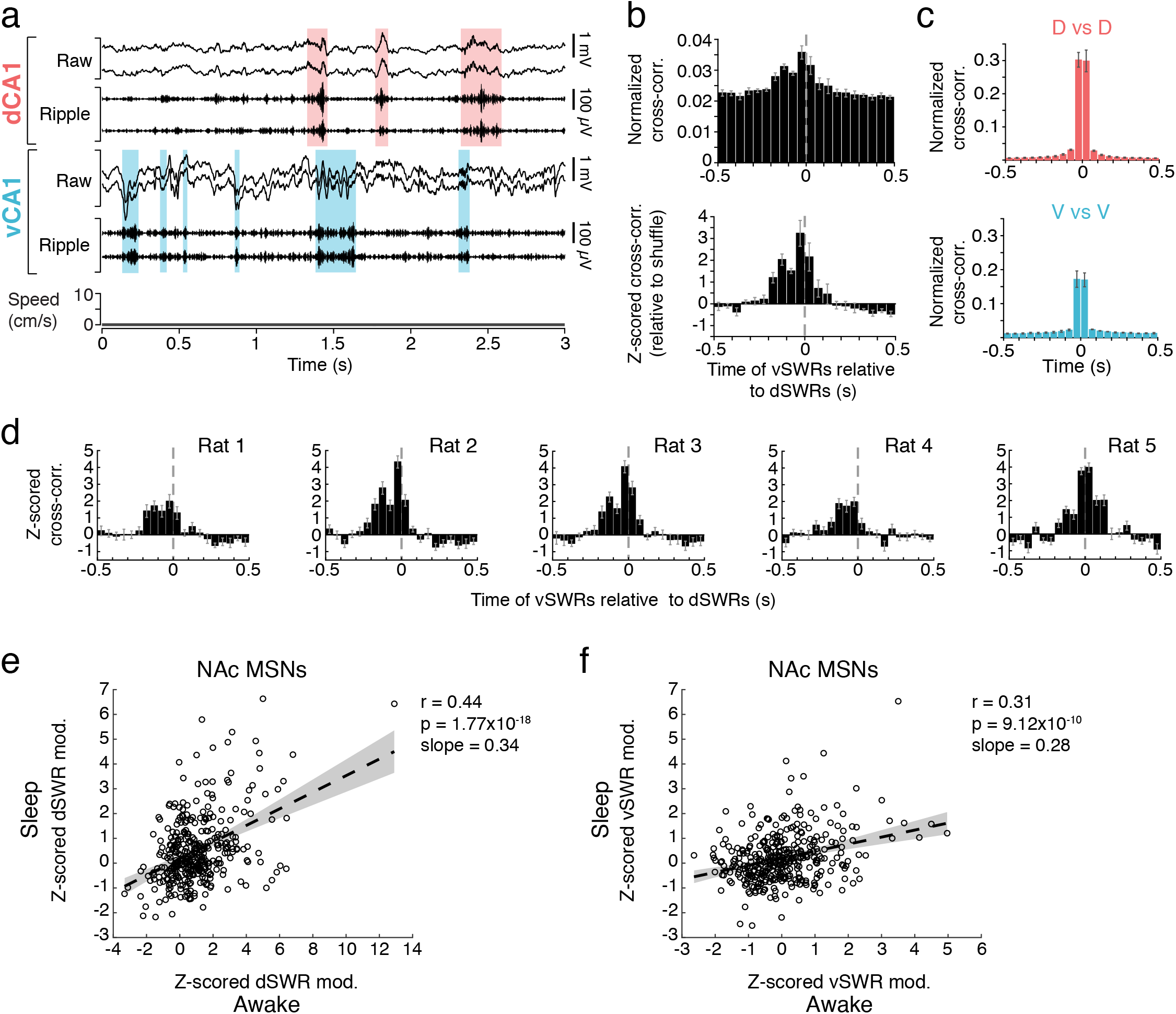
Increased synchrony between dH and vH SWRs during sleep. **a**, Example of dCA1 and vCA1 SWRs during sleep from the same tetrodes and same rat as Fig. 1d (Rat 4). Shaded regions highlight detected dSWRs (pink) and vSWRs (blue). Note the occasional coincidence of dSWRs and vSWRs. **b**, Mean cross-correlation histogram (CCH) for sleep SWRs across animals. Top: normalized by the number of dSWRs. ∼6.7% of dSWRs occur within ±50 ms of a vSWR. Bottom: CCH z-scored relative to shuffled vSWR onset times. Note that synchrony between dSWRs and vSWRs in sleep is greater than expected by chance (>2 z-scores), and vSWRs are most often leading (left of time 0). Error bars indicate s.e.m. (n=5 rats). **c**, High synchrony of sleep SWRs between tetrodes within dH (top, n=5 rats) or vH (bottom, n=3 rats with >1 tetrode in vCA1). Error bars indicate s.e.m. across included animals. **d**, Z-scored CCH (relative to shuffle) of sleep vSWRs and dSWRs in each animal. Error bars indicate s.e.m. across days (n=15-19 days) for each animal. **e**, Z-scored dSWR-modulation of individual NAc MSNs is significantly correlated between awake immobility and sleep dSWRs. Each point represents a single cell, and only cells present in both wake and sleep are included (n=368 cells). Dotted line and shaded regions represent a linear fit with 95% confidence intervals. Pearson’s correlation coefficient (r), p-value and slope are shown in upper right. **f**, Similar to **e**, but for vSWR-modulation of NAc MSNs during awake immobility vs. sleep. Only cells present in both wake and sleep are included (n=368 cells).

